# Integrin conformation-dependent neutrophil slowing obstructs the capillaries of the pre-metastatic lung in a model of breast cancer

**DOI:** 10.1101/2024.03.19.585724

**Authors:** Frédéric Fercoq, Gemma S. Cairns, Marco De Donatis, John B. G. Mackey, Alessia Floerchinger, Amanda McFarlane, Ximena L. Raffo-Iraolagoitia, Declan Whyte, Lindsey W. G. Arnott, Colin Nixon, Robert Wiesheu, Anna Kilbey, Leah Brown, Sarwah Al-Khalidi, Jim C. Norman, Edward W. Roberts, Karen Blyth, Seth B. Coffelt, Leo M. Carlin

## Abstract

Neutrophils are thought to be critical to the process whereby breast cancers establish an immunosuppressive and tumour cell nurturing ‘pre-metastatic’ niche before overt metastasis can be detected. However, the spatial localization of neutrophils and their interaction with other cell types in the lung pre-metastatic niche is not well described. We used a spontaneously metastatic mammary cancer model combined with a multiplexed three- and four-dimensional imaging approach to investigate the behaviour of neutrophils in the pre-metastatic niche. Volume fixed tissue three-dimensional imaging showed that approximately 40% of CD8^+^ T cells are adjacent to neutrophils at this stage. In live tissue, we found neutrophils with impaired intravascular motility congested the capillaries of pre-metastatic lungs potentially obstructing CD8^+^ T cells. Slowed neutrophil transit was dependent on the conformation of β2-integrin and could be recapitulated by treating non-tumour bearing mice with G-CSF, a potent systemic mediator of granulopoiesis. We found a decrease in L-selectin (CD62L) on neutrophils in the lungs of both mammary tumour bearing and G-CSF treated mice. Finally, we observed differential accumulation of intravenously injected micro-beads in the lung, suggestive of transient circulatory dead spaces which were also dependent on β2-integrin inactivation. Overall, our study proposes that integrin-mediated neutrophil congestion of the alveolar capillaries could contribute to the generation of the pulmonary pre-metastatic niche.

## Introduction

Neutrophils have been linked to both pro- and anti-tumour activity based on stage and context^1^ and have been mechanistically implicated in promoting breast cancer metastasis^2–4^. Tumours are known to affect their eventual site of metastasis, even before overt metastasis is detected, forming an environment termed the pre-metastatic niche^5^. Several single cell technologies have been brought to bear on the metastatic and pre-metastatic environment yielding important data on the heterogeneity and plasticity of neutrophils in cancer, but this information often lacks direct spatiotemporal information^6^. Tumours spread to the lung in approximately 30% of metastatic breast cancer patients^7,8^ and these patients have much poorer outcomes^9^. The lung is a unique physiological barrier with a need for carefully balanced regulation between robust immune response and tissue damage. This balance is thought to err on the side of immunosuppression, resolution, and containment to avoid life threatening inflammation^10^. This, along with the extensive narrow vasculature acting as a filter or strainer, makes the lung a key site for metastatic seeding and outgrowth.

Therefore, we probed the behaviour of neutrophils within the lungs of mice bearing spontaneously lung metastatic mammary tumours at a stage before overt lung metastasis could be observed. Using a combination of cleared, fixed, and live *ex vivo* three- and four-dimensional multiplexed immunofluorescence imaging of lung slices we found that the pulmonary capillaries were packed with neutrophils in tumour bearing mice. CD8^+^ T cells within this environment were almost unavoidably in close contact with the neutrophils. Live imaging revealed that the neutrophils moved more slowly than their counterparts in the lungs of healthy mice. This change in neutrophil migratory phenotype was dependent on β2-integrin conformation, a key determinant of neutrophil:endothelium interactions^11^, and was recapitulated by G-CSF treatment in non-tumour bearing mice. Expression of cell surface L-selectin (CD62L), a leukocyte adhesion molecule known to induce β2-integrin activation on neutrophils^12^, was reduced on the surface of lung neutrophils in both mammary tumour bearing, and G-CSF treated mice. We speculate that this aberrant neutrophil intravascular motility is a by-product of phenotypically modified neutrophils from stress granulopoiesis stimulated in part by G-CSF produced by the tumour microenvironment. We suggest that these results are important to consider when targeting T cells or neutrophils in the pre-metastatic niche.

## Results and discussion

### ‘Pre-metastatic’ lung capillaries are neutrophilic

To interrogate the lung pre-metastatic niche we used a murine model of metastatic breast cancer, where either female FVB/N mice, or Catchup^IVM-red^ Ly6G neutrophil reporter mice^13^, were orthotopically transplanted with mammary tumour fragments from *K14-Cre;Trp53^F/F^* (KP) mice^14^ (**Figure 1A**). As in the *K14-Cre;Cdh1^F/F^;Trp53^F/F^* (KEP) transplant model^2^, tumour resection before the mice reach humane endpoint allows them to subsequently develop lung metastases with complete penetrance (**Figure S1**). Therefore, prior to resection, the lungs of these mice recapitulate the pre-overt metastasis microenvironment.

**Figure 1:**
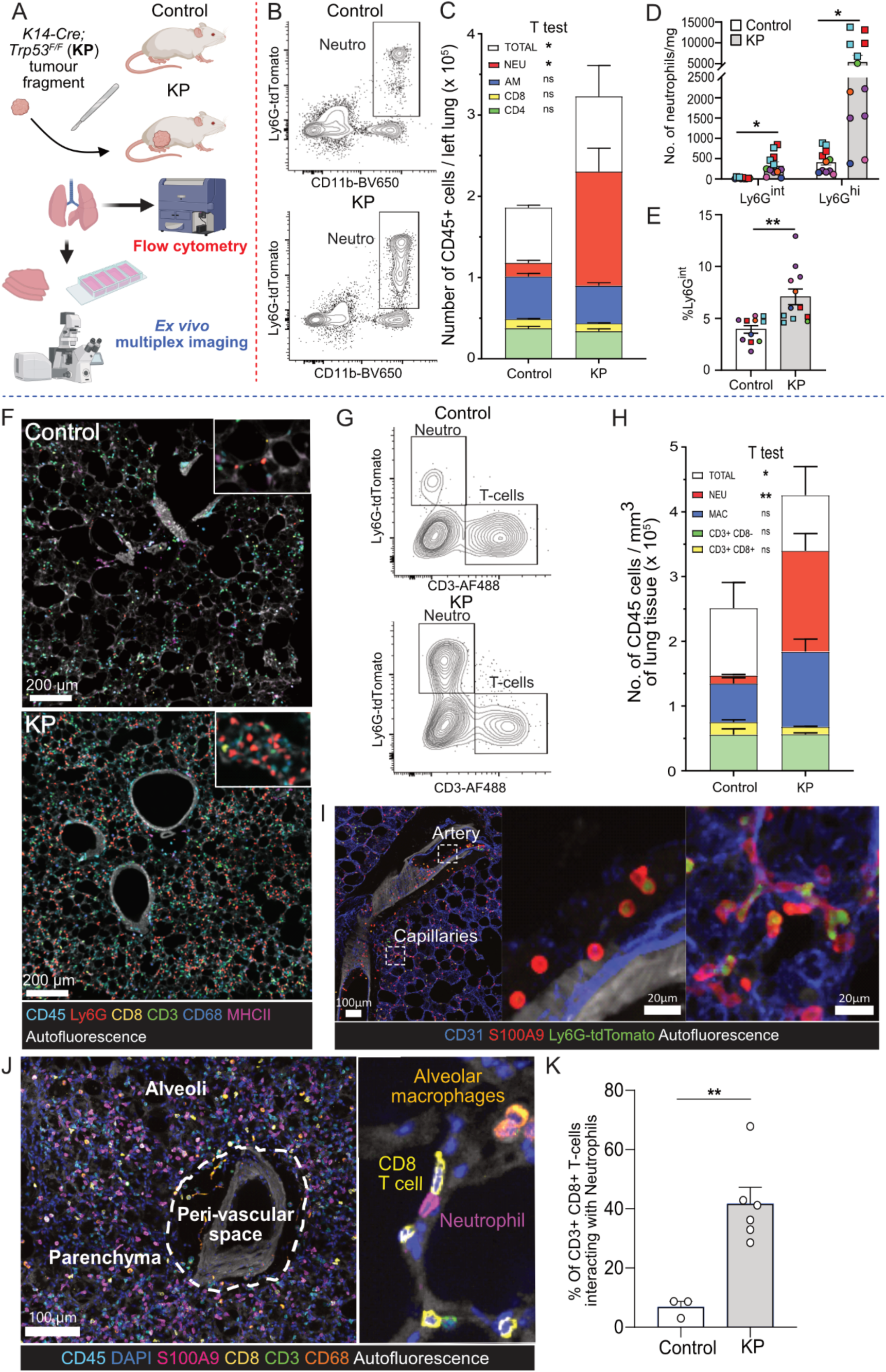
The pre-metastatic lung is congested with neutrophils. (A) Schematic of tumour transplant model and experimental setup. (B-C) Flow cytometry analysis of lungs (using lung immunophenotyping panel) from age-matched non-transplanted control and orthotopic KP 10-15 mm tumour-bearing Catchup^IVM-red^ Ly6G reporter mice. (B) Representative gated plots of CD11b^+^ Ly6G^tdtomato+^ neutrophils, see **Figure S2A** for full gating strategy. (C) Quantification of the different CD45^+^ cell populations. NEU: neutrophils; AM: alveolar Macrophages; CD8: CD8+ T-lymphocytes, CD4: CD4+ T-Lymphocytes. Results are expressed as mean ± SEM of n = 3 control and 6 tumour-bearing Catchup^IVM-red^ mice. Unpaired T-tests were performed for each cell type, *p≤0.05. (D-E) Flow cytometry analysis of Ly6G expression (using neutrophil phenotyping panel) in the lungs. Results are expressed as mean ± SEM of n = 11 control and 12 10-15 mm tumour-bearing mice. Two independent transplant cohorts; squares – cohort one Catchup^IVM-red^ mice, circles – cohort two WT mice. Each colour represents mice which were harvested on the same day when tumours reached endpoint. See **Figure S2B** for gating strategy. (D) Quantification of number of Ly6G^int^ and Ly6G^hi^ neutrophils per mg of lung tissue. A multiple Mann-Whitney test with Holm-Šídák multiple comparisons correction, *p≤0.05. (E) % Ly6G^int^ neutrophils. Unpaired parametric t-test, **p≤0.01. (F-I) Chemically cleared precision-cut lung slices (PCLS) imaged by spectral multiplexed immunofluorescence confocal microscopy as in **Figure S2C**. (F) Maximum intensity projections of spectrally unmixed PCLS images stained with CD3-AF488, CD8-Cy3, CD68-AF594, MHCII-BV421 and a mix of CD45+CD19+CD11c-AF647 in addition to the tdTomato reporter for Ly6G. Autofluorescence was also unmixed and provided tissue architecture information. (G) Leukocytes were segmented, and cell fluorescence intensity was imported in a flow-cytometry analysis software. Figure shows representative plots. (H) Quantification of the different cell populations by histocytometry of fluorescence images. Results, mean ± SEM of n = 3 control and 6 tumour-bearing mice Catchup^IVM-red^ mice. Unpaired T-tests for each cell type, *p≤0.05, **p≤0.01. (I) Fixed PCLS were imaged for neutrophils (S100A9 and Ly6G^tdTomato^) along with CD31 (blue) staining of vascular endothelium. Dashed squares indicate zoomed on an artery area showing round neutrophils or constricted neutrophils in capillaries. (J) Maximum intensity projection of a spectrally unmixed multiplexed confocal microscopy image of a PCLS from a KP tumour bearing FVB/N mouse stained with CD45-Dylight685, DAPI, S100A9-AF700, CD8-Cy3, CD3-AF488 and CD68-AF594. Autofluorescence was also unmixed and provided tissue architecture information. The close-up view of the lung parenchyma shows neutrophil-CD8^+^ T cell interaction in lung capillaries. (K) Quantification of neutrophil-T-cell interactions in PCLS from control and KP-tumour Catchup^IVM-red^ bearing mice. Results as mean ± SEM of n = 3 control and 6 tumour-bearing mice (related to images in figure 1E-G), T-test **p≤0.01. Flow cytometry (B-C) and (D-E) and imaging results (F-I) were obtained from four independent experiments.

In our KP transplant model, we confirmed using flow cytometry, the presence of significant neutrophilia in the pre-metastatic lung of mice with 10-15 mm orthotopic KP mammary tumours (2-6 weeks after transplant, usually week 4) (**Figure 1B, 1C and S2A**), which has been previously reported for the KEP mammary tumour transplant model^2,15^ and other metastatic breast cancer models such as MMTV-PyMT^3^ or 4T1^4,16^. As we previously found in other metastatic models^17^, neutrophilia was accompanied by an increase in both number and percentage of immature Ly6G intermediate (Ly6G^int^) neutrophils (**Figure 1D**, 1E **and S2B**).

Neutrophils and their interaction with T cells have been implicated in breast cancer lung metastasis^2^, however, little is known about the location and interactions of these two cell populations in the pre-metastatic lung. Thus, we used chemically cleared precision-cut lung slices (PCLS) to investigate the immune landscape in the pre-metastatic lung in three dimensions (3D). Spectral multiplexed immunofluorescence confocal microscopy was employed to identify immune cells using six markers simultaneously in large tissue volumes while preserving tissue architecture and unmixing the lungs intrinsic autofluorescence (**Figures 1F and S2C**). Consistent with flow cytometry, histocytometry^18,19^ of 3D multiplexed images demonstrated a substantial accumulation of neutrophils in the lungs of mice with 10-15 mm orthotopic KP mammary tumours, with no statistically significant increase in CD68^+^ macrophage or CD8^+^ T cell abundance (**Figures 1G**, 1H).

Most neutrophils and CD3^+^ T cells were found within the lung vasculature, particularly in pulmonary capillaries and arteries (**Figure 1I and 1J**). We wondered whether the more abundant neutrophils might use the lymphatics to reach the lymph nodes and influence T cells there, but neutrophils were absent from the lymphatic collectors clearly apparent in podoplanin stained sections (**Figure S3A**). Macrophages were predominantly localized in alveoli (alveolar macrophages) or perivascular adventitial spaces (interstitial macrophages) (**Figure 1J)**. In contrast with controls, approximately 40% of CD8^+^ T cells were directly adjacent to neutrophils in tumour bearing mice (**Figure 1J and 1K**) consistent with cell-cell interactions. Notably, the proportion of T cells in contact with neutrophils correlated with the number of lung neutrophils present (Spearman r = 0.8333 p=0.0083) (**Figure S3B**), suggesting that these interactions might be primarily mediated by the significantly increased abundance of neutrophils.

Collectively, these findings indicate that the pre-metastatic pulmonary capillaries are congested with neutrophils. This marked expansion in intravascular neutrophils may facilitate unavoidable contact between neutrophils and CD8^+^ T cells *in vivo*.

### Lung neutrophils do not produce oxidative burst in live PCLS from mammary tumour bearing mice

There is accumulating evidence that upon arrival in the lung, neutrophils acquire unique immunosuppressive capacity to avoid damaging the delicate alveolar tree^4,20,21^. This capacity is thought to be amplified in the context of tumour-driven inflammation^2,4^. Previous work suggested that neutrophils could inhibit cytotoxicity of CD8^+^ T cells in the lung through nitric oxide^2^. We therefore imaged nitric oxide (NO) and superoxide (O2^−^) in live *ex vivo* PCLS to investigate oxidative stress in neutrophils (as depicted in **Figures 2A and S4**). To check the validity of the probes in neutrophils, we stimulated live PCLS with phorbol myristate acetate (PMA) and saw expression of both NO and O2^−^ in neutrophils and surrounding tissue (**Figures S4A and S4D-E**). However, neither NO, nor O2^−^ were detected in lung neutrophils in PCLS from control or KP transplant tumour-bearing mice suggesting that this enhanced neutrophil oxidative burst does not take place in the lungs at this timepoint (**Figures S4B-E**).

**Figure 2:**
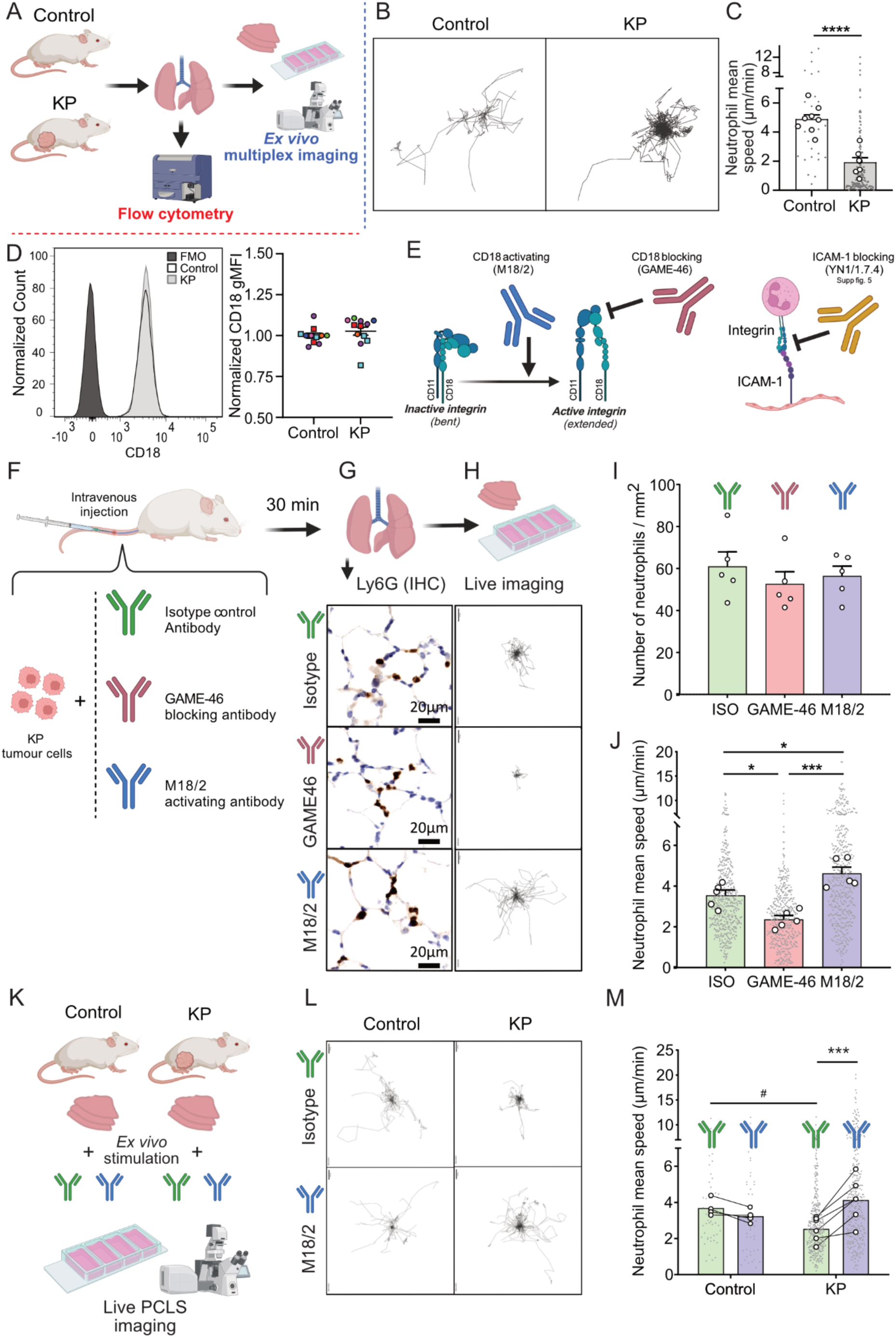
Neutrophils in the pre-metastatic are slow-moving due to integrin inactivation. (A) Schematic of the experiment design. Lungs from age-matched non-transplanted control and 10-15 mm orthotopic KP tumour-bearing FVB/N mice were harvested for live precision-cut lung slices (PCLS) imaging or flow cytometry. (B-C) Neutrophil tracking in live multiplexed confocal microscopy timelapse images of *ex vivo* PCLS from control and KP tumour-bearing mice (see video S1). (B) Representative neutrophil tracking centred plots (spider plot). (C) Quantification of neutrophil mean speed. Grey dots represent the speed of each individual neutrophil (24-148 tracks per group were analysed) while white dots represent the mean speed in each mouse. The mean value per animal were used to perform an unpaired parametric T-test, ****p≤0.0001. Results are expressed as mean ± SEM of n= 8 control and 6 tumour-bearing FVB/N mice. (D) Flow cytometry of CD18 expression on lung neutrophils (using neutrophil phenotyping panel). Left: representative histogram. Fluorescence minus one (FMO) for CD18 also shown. Right: quantification of CD18 geometric mean fluorescence intensity (gMFI). Results are expressed as mean ± SEM of n = 11 control and 12 tumour-bearing mice. Two independent transplant cohorts; squares – cohort one Catchup^IVM-red^ mice, circles – cohort two WT mice. Results normalized to the mean of controls harvested the same day (same coloured points). **Figure S2B** for gating strategy. nonparametric Mann-Whitney test, ns>0.05. (E) Schematic of integrin conformational changes controlling integrin activation and integrin binding to ICAM-1. The M18/2 antibody stabilizes the active conformation, whereas the GAME-46 antibody blocks the CD18 active domain. YN1/1.7.4 antibody blocks ICAM-1 (see **Figure S6**). (F-J) Modulation of neutrophil motility in an intravenous (IV) transplant model of KP cells, using CD18 activating and blocking antibodies. (F) Schematic of the experiment design. Mice were IV injected with KP cells and either isotype control antibody (green), GAME-46 (pink), or M18/2 (blue). 30 minutes later lungs were taken for live *ex vivo* confocal microscopy of PCLS (see **video S2**) and Immunohistochemistry (IHC). (G) Representative IHC from the three treatment groups. (H) Representative neutrophil tracking plots from the three treatment groups. (I) Quantification of the number of neutrophils/mm^2^ in lung IHC. Results expressed as mean ± SEM of 5 mice per group. One-way ANOVA. (J) Quantification of neutrophil speed in live *ex vivo* microscopy. Grey dots represent the speed of individual neutrophils (a total of 482-913 tracks per group were analysed; display was cropped at 512 tracks) while white dots represent the mean neutrophil speed/mouse. Mean value per mouse in ANOVA. (G-J) Results, mean ± SEM of n = 5 mice per group, pool of 5 independent experiments with 1 mouse per group. *p≤0.05, ***p≤0.001. (K-M) *Ex vivo* stimulation of live PCLS from control and KP tumour-bearing mice. (K) Schematic. Live lungs from age-matched controls and 10-15 mm KP-tumour bearing mice, live PCLS stimulated with either isotype control antibody or M18/2 in a chamber slide. (L) Representative neutrophil tracking plots. (M) Quantification of neutrophil mean speed. Grey dots represent the speed of individual neutrophils (a total of 44-371 tracks per group were analysed) while white dots represent the mean neutrophil speed per mouse. Results are expressed as mean ± SEM of n =4 control mice and 6 KP-tumour bearing Catchup^IVM-red^ mice. A Two-way Repeated Measure ANOVA followed with Sídák’s multiple comparisons tests was performed. *** p≤0.001 for differences between isotype and M18/2 antibodies, ^#^p≤0.05 for differences between control and KP tumour bearing mice.

### Neutrophil motility is impaired in a CD18/ICAM-1 dependent manner in the pre-metastatic lung

Further investigation of live PCLS however revealed changes in neutrophil motility in lungs from KP transplant tumour-bearing mice compared to controls. The neutrophils from KP tumour-bearing mice exhibited shorter tracks and reduced mean track speeds compared to those from control mice (**Figures 2B, 2C and Video S1**), while CD8^+^ T cells were mostly immotile in both groups (**Figure S4F**). Therefore, we investigated the mechanisms which control intravascular neutrophil motility in the pre-metastatic lung. Imaging studies have uncovered several examples where the endpoint of leukocyte (including neutrophil): endothelial interactions is not extravasation into the tissue but rather intravascular motility or ‘patrolling’^22–24^. In addition to selectins, neutrophil β2-integrins, Mac-1 (CD11b/CD18; αmβ2) and LFA-1 (CD11a/CD18; αLβ2), which bind to ICAM-1 (CD54) on endothelial cells are thought to mediate neutrophil: endothelial cell interactions^11^. However, due to the narrow diameter of lung capillaries, the typical neutrophil adhesion cascade does not occur. Instead, neutrophils are forced into contact with the endothelium by constriction (**Figures 1I and 1J**). Consequently, it has been postulated that integrins may not be essential for neutrophil recruitment in the lung^25–27^. However, recent studies have suggested a requirement for CD11b (Mac-1) in neutrophil intravascular motility^28^, although it is likely that this regulation extends beyond protein expression, as conformation and intracellular trafficking are known to regulate integrin mediated binding.

Considering the above, we hypothesised that neutrophil intravascular motility in the lungs could be attenuated by integrin inactivation. CD18 (β2), the common beta-chain for both Mac-1 and LFA-1, is expressed by neutrophils at similar levels in both controls and tumour bearing mice (**Figure 2D and S2B**). As integrins undergo conformational changes that govern their adhesion (e.g. to ICAM-1^11^ on the endothelium; **Figure 2E**), we used three monoclonal antibodies to manipulate CD18-ICAM-1 interactions in the lung to investigate this putative mechanism (**Figure 2E**): (1) The CD18 activating antibody (clone M18/2) stabilises the active conformation of CD18, promoting adhesion to ICAM-1^29^; (2) The clone GAME-46 blocks the active domain of CD18, preventing adhesion to ICAM-1^29^; and (3) The ICAM-1 blocking antibody (clone YN1/1.7.4) was used to block ICAM-1. Intravenous injection of tumour cells has been shown to acutely increase neutrophil numbers in lung capillaries^30^, recapitulating that aspect of the pre-metastatic niche in an otherwise unaltered lung environment. We intravenously injected a suspension of KP primary tumour cells plus GAME-46, M18/2, or isotype control antibodies, and lungs were collected 30 minutes later (**Figure 2F**). Subsequently, neutrophils were investigated by immunohistochemistry (IHC) for Ly6G in fixed lung tissue to quantify neutrophil numbers, and in live PCLS to quantify neutrophil speed (**Figures 2G and 2H**). Notably, there was no difference in neutrophil numbers in the lungs across the different treatment groups (**Figures 2G and 2I**). However, CD18 blocking with GAME-46 reduced neutrophil speed, whereas M18/2 increased neutrophil speed compared to isotype control (**Figures 2H and 2J, Video S2**). Likewise, intravenous injection of KP tumour cells with ICAM-1 blocking antibodies or isotype control did not impact neutrophil numbers but blocking ICAM-1 inhibited neutrophil movement in live PCLS (**Figure S5**).

These results suggest that β2-integrin conformation could be a critical modulator of neutrophil motility in the pre-metastatic lung mediated by the primary tumour. Therefore, to investigate whether CD18 conformational stabilisation altered neutrophil speed in the lungs of mice with distant orthotopic KP tumours, we treated live PCLS from control and KP transplant tumour-bearing mice with M18/2 *ex vivo* (**Figure 2K**). Antibody treatment had no significant impact on neutrophil motility in control PCLS; however, M18/2 was able to restore neutrophil crawling speed in PCLS from tumour-bearing mice (**Figures 2L**, 2M **and S6A**). This suggests that tumour-driven neutrophil motility impairment could be mediated in part by conformational inactivation of CD18.

### G-CSF recapitulates decreased lung neutrophil motility

We next investigated the potential mediators of CD18 conformational change. It is known that breast cancers induce neutrophilia through G-CSF^2–4^. We therefore wanted to probe whether G-CSF could drive the neutrophil slowing we had observed in the tumour-bearing mice in the absence of a tumour. Mice were administered a single 5 µg dose of G-CSF intraperitoneally, and the lungs were collected 24 hours later to analyse neutrophil motility in *ex vivo* live PCLS, and neutrophil abundance by IHC and flow cytometry (**Figure 3A**). In live PCLS, neutrophil behaviour recapitulated that observed in tumour-bearing mice (**Figures 3B, 3C and 2M**). Notably, neutrophil speed was reduced in G-CSF-treated mice, which could be rescued with M18/2 as before (**Figures 3B, 3C and S6B**). As expected, lung neutrophil numbers were substantially significantly increased following G-CSF treatment (**Figures 3D, 3E and 3F**), with a trend for increased absolute number (p=0.07) and significantly increased (p≤0.001) percentage of Ly6G^int^ neutrophils (**Figure 3F and 3G**). In line with the appearance of immature neutrophils, CD101, which has been used as a neutrophil maturity marker in other cancer models^31^, decreased by approximately 50% with G-CSF treatment (**Figures 3H and 3I**) and more moderately by approximately 20% in the tumour-bearing mice (**Figures 3J and 3K**).

**Figure 3:**
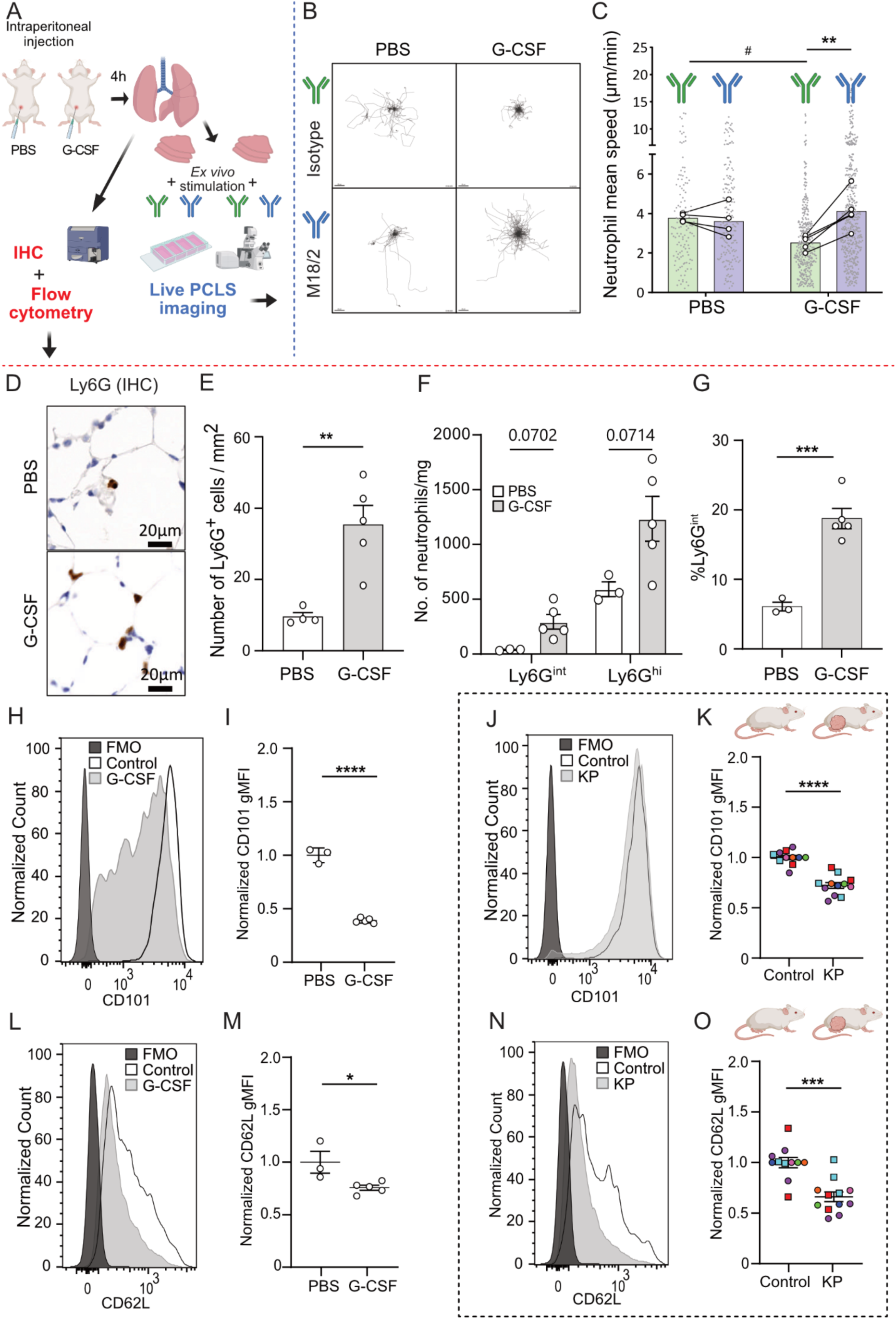
G-CSF induces mobility inhibition by impairing CD18 activation. (A) Schematic of experiment design. Mice were IP injected with either PBS or 5 µg G-CSF. 24 hours later lungs were taken for IHC, flow cytometry and/or live *ex vivo* multiplexed confocal microscopy of PCLS stimulated with either isotype control or M18/2. (B) Representative neutrophil tracking plots of live PCLS from PBS and G-CSF treated FVB/N mice, stimulated *ex vivo* with either isotype antibodies or M18/2. (C) Quantification of mean neutrophil speed from live multiplexed confocal microscopy. Grey dots represent the speed of individual neutrophils (a total of 128-482 tracks per group were analysed) while white dots represent the mean speed per mouse. A Two-way Repeated Measure ANOVA followed with Sídák’s multiple comparisons test was performed on the mean values. Results are expressed as mean ± SEM of n = 4 PBS control and 5 G-CSF treated FVB/N mice, ** p≤0.01 for differences between isotype and M18/2 antibodies, ^#^p≤0.05 for differences between control and G-CSF treated mice. (D) Representative Ly6G IHC images from PBS and G-CSF treated FVB/N mice. (E) Quantification of number of Ly6G^+^ cells (neutrophils)/mm^2^ of lung tissue in IHC images. Results are expressed as mean ± SEM of n = 4 PBS control and 5 G-CSF treated mice. An unpaired parametric T-test was performed. **p≤0.01. (F-O) Flow cytometry of lung neutrophils (using neutrophil phenotyping panel), see **Figure S2B** for gating strategy. (F-I, L-M) Mean ± SEM of n = 3 PBS control and 5 G-CSF treated Catchup^IVM-red^ mice. (J-K, N-O) Comparable flow cytometry of lung neutrophils in KP transplant mice (black dashed box). Mean ± SEM of n = 11 control and 12 tumour-bearing mice. Two independent transplant cohorts; squares – cohort one are Catchup^IVM-red^ mice, circles – cohort two are FVB/N WT mice. (F) No. Ly6G^int^ immature and Ly6G^hi^ mature neutrophils/mg lung tissue. Mann-Whitney with Holm-Šídák multiple comparisons correction. (G) % of Ly6G^int^ neutrophils. T-test ***p≤0.001. (H) Representative CD101 histogram. (I) CD101 gMFI normalized to mean of controls. T-test ****p≤0.0001. (J) Representative CD101 histogram. (K) CD101 gMFI normalized to the mean of the controls harvested on the same day (same coloured points). T-test ****p≤0.0001. (L) Representative CD62L histogram. (M) CD62L gMFI normalized to mean of controls. T-test *p≤0.05. (N) Representative CD62L histogram. (O) CD62L gMFI normalized to the mean of the controls harvested on the same day (same coloured points). T-test, ***p≤0.001.

### Reduced CD62L cell surface expression in lung neutrophils from G-CSF treated and KP mammary tumour bearing mice

Interestingly, further immunophenotyping of lung neutrophils by flow cytometry (**Figure S2B**) revealed that the level of cell surface CD62L was reduced in both G-CSF treated (**Figure 3L and 3M**) and KP mammary tumour-bearing mice (**Figure 3N and 3O**). Interestingly, CD62L^dim^ neutrophils have been connected with breast cancer lung metastasis^32,33^. As noted in the introduction, CD62L has been implicated in β2-integrin activation on neutrophils^12^, therefore, there may be a mechanistic link between CD62L (either by reduced expression or shedding) and β2-integrin conformation which results in neutrophil impaired motility in our model. These and the findings above are consistent with the reduction in pro-metastatic neutrophils previously observed upon blocking G-CSF in mammary tumour bearing mice^2^.

### Slow moving neutrophils alter blood flow

Finally, we aimed to determine the consequences of neutrophilia and neutrophil speed impairment in lung vasculature. In the context of the work above, we hypothesized that neutrophils could (1) favour the accumulation of circulating tumour cells arriving in lung capillaries as neutrophils have been shown to interact with tumour cells in liver sinusoids and favour their arrest^34,35^; and (2) neutrophils could affect normal blood circulation, as it has been shown that neutrophils can obstruct capillaries in the brain^36,37^ and lung^38^, impairing blood flow and creating transient circulatory dead spaces.

Therefore, we intravenously administered fluorescent beads with a diameter of 10 µm (larger than typical lung capillary size) and 4 µm (slightly smaller than a red blood cell) to control and KP tumour-bearing mice to both ‘model’ tumour cells and to have a readout of lung blood volume respectively (**Figure 4A**). In addition, mice were treated with either M18/2 or isotype control antibodies to determine whether restoring neutrophil motility affects bead retention in the lung (**Figure 4A**). IHC staining for Ly6G on lung tissue did not indicate a significant difference in neutrophil accumulation in the lungs of tumour-bearing mice treated with isotype control or M18/2 antibody (**Figures 4B and 4C**), in line with what we had observed in the IV-injection model (**Figures 2F and 2I**). 3D fluorescence imaging of fixed PCLS revealed that the retention of 10 µm beads remained unaffected by the presence of primary tumours or antibody treatment, likely due to physical entrapment in the capillaries (**Figures 4D and 4E**). However, significantly fewer 4 µm beads were found in PCLS from tumour-bearing mice treated with isotype control when compared with control mice, supporting the idea of transient formation of circulatory dead spaces in the lung reducing total blood volume (**Figure 4F**). Interestingly, treatment with M18/2 antibody was able to restore a similar bead count to that from control mice, suggesting that restoring neutrophil motility also restores blood circulation (**Figure 4F**). The formation of transient poorly perfused spaces could prevent efficient immune surveillance and protect the tumour cells, supporting a true ‘pre-metastatic niche’.

**Figure 4.**
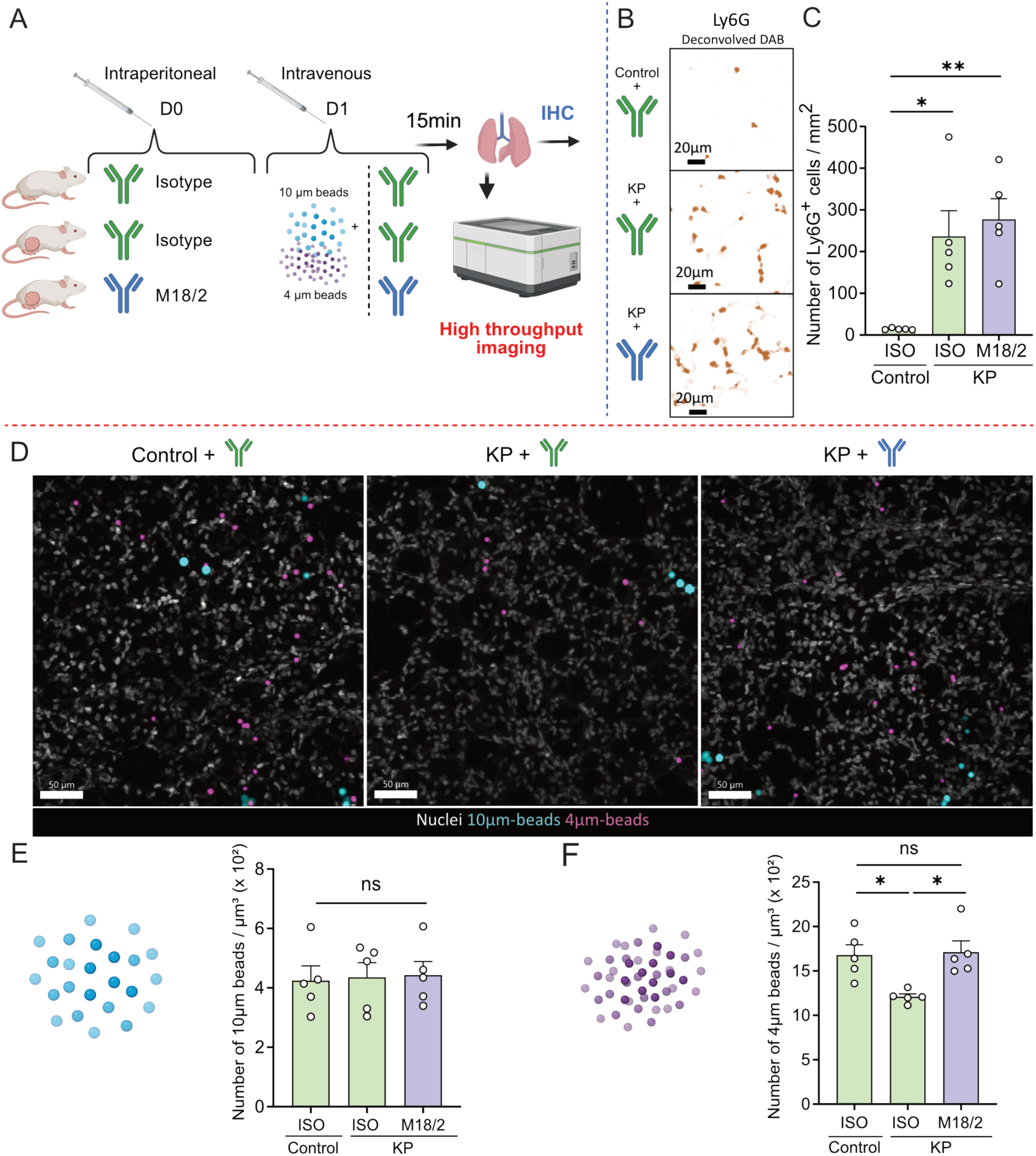
Slowed neutrophils in the pre-metastatic lung restricts blood flow. (A) Schematic of experiment design. 4- and 10-µm fluorescent beads were intravenously injected to control and KP tumour-bearing FVB/N mice along with either M18/2 or isotype control antibodies. Lungs were there harvested for (B-C) Ly6G IHC quantification and (D-F) high throughput confocal fluorescence imaging of fixed PCLS. (B) Representative IHC images from the three different experiment groups. (C) Quantification of the number of Ly6G^+^ cells per mm^2^ of lung tissue. Results are expressed as mean ± SEM of n = 5 mice per group. A one-way ANOVA with Šídák’s multiple comparison test was carried out (*p≤0.05, and **p≤0.01). (D-F) High throughput confocal fluorescence imaging of fixed PCLS to quantify number of beads in the slices. (D) Representative individual field of views for the three experiment groups where DNA was labelled with DR. (E) Quantification of the number of 10µm beads per µm^3^. (F) Quantification of the number of 4µm beads per µm^3^. Results are expressed as mean ± SEM of n = 5 mice per group (same mice as in B-C). A one-way ANOVA with Šídák’s multiple comparison tests was carried out (*p≤0.05).

Taken together, our results suggest that when primary mammary tumours induce substantial systemic neutrophilia, this aberrant neutrophil population fail to activate the β2-integrins that would allow normal intravascular motility, causing congestion and impairment of local blood circulation. That this can be recapitulated by treatment of G-CSF to non-tumour bearing mice suggests a role for this potent mediator of neutrophil homeostasis, known to be increased in the tumour microenvironment of mammary tumours^15^.

In the context of breast cancer metastasis, impaired intravascular neutrophil motility, congestion and reduced local circulatory flux could offer niches inaccessible to anti-tumour immune cells and promote tumour cell seeding indirectly. Additionally, CD11b/CD18 has also been implicated in neutrophil antibody-dependent cellular cytotoxicity (ADCC) of tumour cells^39^. Inactivation of neutrophil integrins could therefore provide another mechanism of immune escape mediated by the primary tumour to favour metastasis at distant sites.

## Materials & Methods

### Mice

Either female FVB/N mice purchased from Charles River Laboratories (acclimatised for at least 1 week in our mouse facility), or female Catchup^IVM-red^ mice^13^ (identified as *Ly6G^Cre-tom/+^ ROSA26^tdtom/+^*, backcrossed onto a FVB/N background for >10 generations and bred in-house) were used for experiments when they were 8-15 weeks old. Age matched mice were randomly allocated to control and transplant/treatment groups. How specific cohorts were analysed is described in the appropriate methods sections and figure legends.

Mice were housed (as in ^17^) at the CRUK Scotland Institute, UK in specific pathogen-free conditions with a 12 hr light/dark cycle and at a controlled climate of 19-22 °C and 45-65 % humidity. Mice also had access to food and water ad libitum. Mouse procedures were performed in accordance with the UK Animal (Scientific Procedures) Act 1986, approved by the local animal welfare (AWERB) committee, and conducted under UK Home Office licences (PP6345023, P72BA642F and PP0826467) at the CRUK Scotland Institute.

### Tumour models

#### Pre-metastatic lung induction

To induce a pre-metastatic lung, 1 mm^3^ mammary tumour fragments obtained from *K14-Cre;Trp53^F/F^* (KP) mice (maintained on an FVB/N background)^14^ were orthotopically transplanted into the 4^th^ mammary fat pad of 10-12 week old female recipient FVB/N or CatchupIVM-red mice using anaesthesia and analgesia. The tumour pieces were allowed to develop into a primary mammary tumour of 10-15 mm in any direction measured three times a week using a calliper and mice were humanely killed usually within a month after transplant. Control mice did not undergo transplantation but were kept in the same conditions.

#### IV injection models

For intravenous injection, sub-confluent KP cells, initially obtained from a mammary KP tumour as described before^40^, were washed, trypsinized, centrifuged (300 g, 5 min), resuspended in medium (RPMI) and passed through a 70 µm cell strainer (Greiner Bio-One). The cells were counted using the CellDrop cell counter (DeNovix), diluted to a maximal concentration of 10^6^ cells per mL if necessary and stained with CellTrace (Invitrogen) according to the manufacturer’s protocol (Invitrogen). Briefly, cells were incubated with 1 μM CellTrace yellow or far-red in PBS for 20 min at 37 °C. The suspension was then diluted fivefold with culture medium (RPMI) and incubated for 5 min to remove free dye in the solution. Finally, the suspension was centrifuged (300 g, 5 min) and resuspended in PBS before mice were injected intravenously with 100 µL of suspension of 0.5x10^6^ cells and 10 µg of monoclonal antibodies. The following low endotoxin, azide-free purified antibodies were used: isotype control (clone RTK2758, Biolegend), ICAM-1 blocking antibody (clone YN1/1.7.4, Biolegend), CD18-activating antibody (clone M18/2, Biolegend), CD18-blocking antibody (clone GAME-46, BD biosciences).

### G-CSF treatment

Female Catchup^IVM-red^ and FVB/N mice aged 8-15 weeks were treated with 5 µg mouse recombinant G-CSF (STEMCELL Technologies) by intraperitoneal injection (i.p.) 24 hours before harvesting the stated tissues.

### Flow cytometry

Lungs were excised and then minced with scissors before incubation in RPMI (ThermoFisher) with 10% foetal bovine serum (FBS, ThermoFisher) at 37 °C for 20 min with agitation. Post-incubation, lungs were mashed through a 70 μm filter.

Single cell suspensions were stained with Live/Dead stain (LIVE/DEAD Fixable Near-IR, Yellow or Green, Biolegend) and Fc-Receptor block was performed (clone 93, Biolegend). Cell suspensions were incubated with directly conjugated fluorescent antibodies in Brilliant Stain Buffer (BD Horizon) at 4 °C for 30 min.

Count beads (CountBright, Life Technologies) were used to determine cell numbers. Compensation was performed using single stained beads (Ultracomp eBeads, ThermoFisher). Acquisition was performed on a BDFortessa using FacsDiva software (BD Biosciences) and data were analysed using FlowJo software (BD).

The following antibodies were used:

Lung immunophenotyping panel: (all 1:200): CD8-AF488 (clone 53-6.7, Biolegend), CD45-BV711 (clone 30-F11, Biolegend), CD11b-BV650 (clone M1/70 Biolegend), SiglecF-BV605 (clone E50-2440 BD Biosciences), F4/80-BV510 (clone BM8, Biolegend), B220-BV421, Ly6G-BUV395 (clone 1A8, BD Biosciences), CD3-AF657 (clone 17A2, Biolegend), CD4-PE/Cy7 (clone RM4-5, Biolegend).

Neutrophil phenotyping panel: Ter119-FITC (clone TER-119 Biolegend 1:200), NKp46-FITC (clone 29A1.4 Biolegend 1:200), CD115-AF488 (clone AFS98 Biolegend 1:200), CD3-FITC (clone 17A2 Biolegend 1:200), CD19-FITC (clone 6D5 Biolegend 1:200), CD45-BV650 (clone 30-F11 Biolegend 1:200), CD11b-BV605 (clone M1/70 Biolegend 1:400), Ly6G-BV510 (clone 1A8 Biolegend 1:200), Siglec-F-APC-Cy7 (clone E50-2440 BD Biosciences 1:200), CD18-AF647 (clone M18/2 Biolegend 1:200), CD62L-BUV395 (clone MEL-14 BD Biosciences 1:200), CD101-PE-Cyanine7 (clone Moushi101 Invitrogen 1:200).

### Fixed PCLS imaging

Precision-cut lung slices (PCLS) were prepared as previously described ^41–43^. Mice were humanely killed by overdose of anaesthetic (i.p. injection of sodium-pentobarbital) followed by permanent cessation of the circulation by cutting the femoral artery. A blunted 22G needle was then inserted via a tracheostomy and used to inflate the lungs slowly but continuously with 1 mL of 2% low-melting point agarose (ThermoFisher) prepared in PBS. When flow cytometry of lung tissue was to be performed on the same animals, the left lung was tied off with a suture before lung inflation. Using suture thread, the trachea was then tied to stop agarose leakage before excising lungs, heart and trachea together and placing them in ice-cold PBS. They were then subsequently fixed in 4% formaldehyde (v/v) (FA) in PBS (ThermoFisher) for two hours or overnight. 300 μm lung tissue sections were sliced using a vibrating microtome (5100mz; Campden Instruments) and stainless-steel blades (Model 7550-1-SS; Campden Instruments) with the highest oscillation amplitude, maximal speed, and an advance of 1 mm/sec. Lung sections were stored in PBS/1% (w/v) BSA (Merck)/0.05% (v/v) Sodium Azide (GBiosciences) (PBA) at 4°C. For permeabilization and blocking, samples were incubated in 300 μL permeabilization buffer (PBS, 1% BSA, 0.05% Azide, 10% NGS, 0.3% Triton X 100) for at least one hour. Slices were rinsed with wash buffer (PBS, 1% BSA, 0.05% Azide, 0.1% Triton X 100) before staining with primary antibodies diluted in antibody buffer (PBS, 1% BSA, 0.05% Azide, 10% NGS, 0.1% Triton X 100). Incubation at room temperature in the dark was allowed for at least 3 hours or overnight before washing three times with wash buffer at 37 °C for at least 20 min each. Directly conjugated antibodies, secondary antibodies, and nuclear stains (DAPI or Draq5, 1:1000) diluted in antibody buffer were incubated with the samples for at least 3 h in the dark. If primary and directly conjugated antibodies were derived from the same species (i.e. rat), they were not stained simultaneously. In this case, the slices were washed as above after incubation with the secondary antibody and free antigen binding sites were blocked with appropriate isotype to avoid binding of the secondary antibody to the subsequently used directly conjugated antibodies. Slices were washed twice as above and once with PBS before fixation with 4% FA for 5 mins. FA was removed with two PBS washes before tissue clearing was performed. Clearing with Ce3D including the preparation of the clearing solution has previously been described in detail^19,44^. Briefly, N-methylacetamide was melted and diluted in PBS to prepare a 40% (v/v) stock solution. 86% Histodenz (w/v) was added and allowed to fully dissolve by incubation at 37 °C in a shaking incubator. 0.1% (v/v) Triton X 100 and 0.5% (v/v) 1 thioglycerol were added. Slices were immersed in Ce3D and incubated at room temperature until sufficient clearing was observed. The tissue sections were mounted on microscope slides (VWR) in Ce3D using Frame Seal incubation chambers (Bio Rad) and kept at 4°C in the dark after coverslipping (thickness no. 1.5; VWR). Slices were imaged on a Zeiss LSM 880 confocal microscope using a 32 channel Gallium arsenide phosphide (GaAsP) spectral detector with plan Apo objective lenses (20X/0,8 NA - Air, Carl Zeiss). Samples were excited simultaneously with 405, 488, 561 and 633 laser lines and signal was collected onto the arrayed GaAsp detector in lambda mode with a resolution of 8.9 nm over the visible spectrum. Using reference spectra acquired from either unstained tissue for autofluorescence or single stained tissue for each antibody listed below, the images were then spectrally unmixed using Zen software (Carl Zeiss).

The following primary antibodies were used: Hamster anti-mouse CD31 (1:400, clone 2H8, Invitrogen), rat anti-mouse CD8 (1:200, 14-0808-82, ebioscience), S100A9 (1:1000, EPR22332-75, Abcam).

The following secondary antibodies were used: Goat Anti-Armenian Hamster polyclonal IgG conjugated with AF488 (1:400, Jackson Immunoresearch), Goat Anti-Rat IgG conjugated with Cy3 (1:400, Jackson Immunoresearch), Goat anti-Rabbit IgG conjugated with AF700 (1:400, Invitrogen). Antibodies with minimal Cross Reactivity to human, bovine, horse, mouse and rabbit serum proteins were preferred.

The following conjugated antibodies were used:

Anti-mouse I-A/I-E conjugated with BV421 (1:200, clone M5/114.15.2, Biolegend), anti-mouse Podoplanin conjugated with AF488 (1:200, clone PMab-1, Biolegend), anti-mouse CD3 conjugated with AF488 (1:200, clone 17A2, Biolegend), anti-mouse CD68 conjugated with AF594 (1:400, clone FA-11, Biolegend)

### Cell segmentation and histocytometry analysis

For histocytometry analysis, the method previously published by Li et al.^19^ was adapted to the specific needs of the experiment. Gaussian filtering in Imaris was applied on the channels of interest representing specific cell types. Summing was used to generate a merged channel with all stained cell types using the Channel Arithmetic function in Imaris. Multiplier factors served to normalize the channels to one another and obtain similar intensities for all markers in the composite channel. Surface objects were created with the respective module in Imaris. The surface detail in the smooth function was set to 1.4 μm. Background correction was enabled to allow segmentation based on local contrast in areas with differing background intensities such as bronchi with high autofluorescence. A threshold was set to incorporate all cell derived signals while excluding unspecific background. The split touching objects option was enabled and set to 8 μm, i.e. the approximate mean size of the observed cell types. Seed points were filtered to only include events with sufficient signal intensity. After verifying an adequate representation of the cells of interest by the created surface objects, the statistics for these objects were exported into CSV files.

Relevant parameters were combined into one Excel file applying the RDBMerge plugin (http://www.rondebruin.nl/win/addins/rdbmerge.htm) for automation of the process. In our experiments, mean and summed channel intensity, volume, area, and position are the crucial parameters. These data were saved in a CSV file which can subsequently be opened in FlowJo for phenotypic gating and quantification of different cell populations like typical flow cytometry analysis.

Interactions between stained CD8^+^ T cells and Ly6G^+^ Neutrophils were quantified based on the distance between one cell and the nearest cell of the other cell type. To achieve this, spots and surface objects were generated in Imaris for both cell types respectively and the ‘Find spots close to surface’ XTension was used to define the subset of cells that interact with the other cell type, i.e. spots that are within the set threshold distance of 8 μm from a surface object.

### Live PCLS imaging

All mice used for neutrophil dynamics analysis were humanely killed within the same period (between 09.00-10.30) to minimize circadian-induced changes between samples. Where cohorts needed to be split over several imaging sessions, each imaging session was carried out with at least one age-matched control. Live PCLS procedure was performed as in ^41^. Lungs were inflated with low melting agarose point as above. Excised lungs were placed in ice-cold PBS. Lungs were sliced into 300μm thick sections on a vibratome and stained with directly conjugated antibodies, Hoechst and probes for reactive superoxide and nitric oxide (ROS-ID® ROS/RNS detection kit, Enzo Life Sciences) in complete medium (phenol-red free DMEM substituted with 10% FBS) for 20 minutes at 37°C. For *ex vivo* integrins activation experiments, isotype control (clone RTK2758, Biolegend) or CD18-activating (clone M18/2, Biolegend) antibodies were added to the solution at 1:50 dilution.

The following antibodies were used (all 1:200): Anti-mouse CD31-BV421 (clone 390, Biolegend), anti-mouse CD8-AF488 (clone 53-6.7, Biolegend), anti-mouse Ly6G-DyLight550 (clone 1A8, Novus Biologicals), anti-mouse CD11c-AF594 (clone N418, Biolegend), anti-mouse CD45 Spark NIR685 (clone 30-F11, Biolegend).

Slices were imaged on a Zeiss LSM880 confocal microscope using multiwell incubation chambers (Ibidi µ-Slide 8-well or Nunc Lab-Tek II 8-well Chamber Slide System) at 37°C with 5% CO2. Lung slices were imaged for 6 to 40min with z-stacks of 22-35µm. Acquisition was performed with a 32 channel Gallium arsenide phosphide (GaAsP) spectral detector plan Apo objective lenses (20X/0,8 NA - Air, Carl Zeiss). Samples were excited simultaneously with 405, 488, 561 and 633 laser lines and signal was collected onto a linear array of the 32 GaAsp detectors in lambda mode with a resolution of 8.9 nm over the visible spectrum. Spectral images were then unmixed with Zen software (Carl Zeiss) using reference spectra acquired from unstained tissues (tissue autofluorescence) or slides containing single fluorophores.

### 4D image analysis

Timelapse image analysis and visualization was performed using Imaris (Bitplane). Neutrophils and CD8^+^ T cells were segmented and tracked using the ‘surface’ tool using either Ly6G fluorescence intensity (neutrophils) or CD8 (T-cells). All surfaces and tracks were checked manually to avoid any false detections. Oxidative stress was measured with superoxide/nitric oxide mean cell fluorescence and cell behaviour was determined using the mean track speed.

### Immunohistochemistry (IHC)

Lungs were fixed in 4% buffered formalin overnight. Fixative was changed 24 h post-fixation for a further 24 h. Thereafter, lungs were dehydrated in increasing concentrations of 70% to 100% ethanol, and then placed in xylene before paraffin embedding. Four-micron-thick serial sections were prepared as previously published^43^. Immunostaining for neutrophils was performed at the CRUK Scotland Institute’s Histology Facility. Antigen retrieval was performed using ER2 retrieval buffer (20 min, 95 °C). Sections were then stained for Ly6G (clone 1A8) at a previously optimized dilution (1:60000) for 30 min at room temperature with the autostainer Bond Rx (Leica Microsystems GmbH, Wetzlar, Germany). The IHC sections were visualised with 3,3’Diaminobenzidine (DAB). Stained sections were then digitally scanned using a Leica Aperio AT2 slide scanner at ×20 magnification. Digital slides were analysed with QuPath^45^. Lung sections were annotated using ‘pixel classification’ and DAB positive cells within the annotated areas were detected using the “positive cell detection” tool. Results were then expressed as the number of positive cells/mm^2^ of lung tissue.

### Fluorescent beads retention in the lung

Once outgrown tumours reached 12mm in any direction, mice were injected intraperitoneally with 20µg of CD18 activating antibody (clone M18/2, Biolegend) or Isotype control (clone RTK2758, Biolegend) in 100µL PBS. 24h later, mice were injected intravenously with a 100µL mixture containing 10µg of CD18 activating antibody or isotype control, 500 000 blue-fluorescent Polystyrene Microspheres, diameter 10 µm (FluoSpheres, Thermofisher) and 20uL of a solution (2% solids) of yellow/green-fluorescent Sulfate Microspheres, diameter 4.0 µm (FluoSpheres, Thermofisher). Mice were culled for lung collection 15 minutes after injection and lung were processed for fixed PCLS imaging (see above). Cleared PCLS were imaged using an Opera Phenix (Perkin Elmer) high throughput spinning disk microscope. Images were analysed using Imaris (Bitplane) to measure the number of beads / mm^3^ in the lung sections.

### Statistics

Representation and data analysis were performed with Prism 9.0 and 10.0 software (GraphPad Inc.). The choice of statistical analysis was based on sample size and distribution of samples. Normality was assessed via the Shapiro–Wilk test. Data from independent experiments were pooled when possible. Where normality could be determined (1) parametric t-tests were used to compare two groups; (2) One-way analysis of variance (ANOVA) was used to compare three groups according to one variable (antibody treatment); (3) Two-way ANOVA, followed with Sídák’s multiple comparisons tests was performed to analyse the influence of two independent variables (tumour presence and/or antibody treatment). When normality could not be determined, or data were not normally distributed, multiple Mann-Whitney test was performed with multiple comparisons correction using the Holmes-Sídák to compare multiple groups. Specific tests used in each case are stated in the figure legends. Means +/-standard error of the mean are indicated were possible. Specific numbers of animals can be found in corresponding figure legends.

## Acknowledgments

We are grateful to Catherine Winchester (CRUK Scotland Institute) for critical reading and checking of the manuscript. We are grateful for support from all the services and resources at the CRUK Scotland Institute with particular thanks to the Biological Services Unit, the Beatson Advanced Imaging Resource (BAIR), the Flow Cytometry Facility and the Histology Facility. We also thank the MNHN light microscopy facility (CeMIM, Centre de Microscopie et d’IMagerie numérique, MNHN Paris) for providing access to Confocal scanning laser microscopes. We thank Jos Jonkers (Netherlands Cancer Institute) for the KP mammary cancer mouse model. Illustrations were created with BioRender.com.

## Data availability

Upon acceptance, all data will be made available by the corresponding author on request.

## Competing interests

LMC has consulted for Ono Pharmaceuticals UK, JBGM has subsequently become an employee of AstraZeneca, AJM has subsequently become an employee of Antibody Analytics

## Funding

FF, GSC, SBC & LMC were supported by Breast Cancer Now grant 2019DecPR1424 and CRUK Scotland Centre core - CTRQQR-2021\100006

LMC, FF, GSC, MDD, JBGM, XLRI, AM, LWGA were supported by CRUK SI core programme to LMC A23983 and CRUK SI core funding to the CRUK Scotland InsVtute A31287

LMC - DRCRPG-Nov22/100007

JCN, DW - CRUK Core A28291

CN - CRUK SI core A31287

LB – Wellcome Trust Biomedical VacaVon Scholarship 220281/Z/20/Z

EWR - CRUK SI core A1920

KB - CRUK SI Core A29799 CRUK SI core A31287

## Author contributions

<u>**Conceptualization**</u> – FF, GSC, SBC, LMC

<u>**Formal analysis**</u> – FF, GSC

<u>**Funding acquisition**</u> – JCN, EWR, KB, SBC, LMC

<u>**Investigation**</u> – FF, GSC, MDD, JBGM, AF, AM, XLR, DW, LWGA, CN, RW, AK, LB, SA-K

<u>**Methodology**</u> – FF, KB, SC

<u>**Project administration**</u> – LMC

<u>**Resources**</u> – RW, AK, SBC

<u>**Supervision**</u> – JCN, EWR, KB, SBC, LMC

<u>**Validation**</u> – FF, GSC, DW

<u>**Visualization**</u> – FF, GSC

<u>**Writing – original draft**</u> – FF, GSC, LMC.

## Supplementary figures and legends

**Video 1: Neutrophil behaviour and free radical production in live PCLS.**

Lungs from 10-15 mm KP tumour bearing mice were harvested for live *ex vivo* multiplexed confocal microscopy of PCLS to analyse their behaviour and free radical production (Reactive Oxygen Species – ROS and Reactive Nitrogen Species – RNS). Slices were stained for DNA (Hoechst, blue), vasculature (CD31, grey) and neutrophils (Ly6G, red) as well as probes for ROS (orange) and RNS (yellow). 5 minutes timelapse series were acquired and neutrophils were tracked using Imaris software.

**Video 2: CD18 modulates lung neutrophil motility in an intravenous transplant model of KP cells.**

Catchup^IVM-red^ mice were IV injected with KP cells stained with cellTrace dyes (yellow) and either isotype control antibody, CD18 blocking antibody (GAME-46) or CD18 activating antibody (M18/2). 30 minutes later lungs were taken for live *ex vivo* confocal microscopy of PCLS. Slices were stained for DNA (Hoechst, blue), vasculature (CD31, grey), neutrophils (Ly6G, purple) and CD45 (cyan). 40 minutes timelapse series were acquired, and neutrophils were tracked using Imaris software.

**Video 3: ICAM-1 modulates lung neutrophil motility in an intravenous transplant model of KP cells.**

Catchup^IVM-red^ mice were IV injected with KP cells stained with cellTrace dyes (yellow) and either isotype control antibody or ICAM-1 blocking antibody. 30 minutes later lungs were taken for live *ex vivo* confocal microscopy of PCLS. Slices were stained for DNA (Hoechst, blue), vasculature (CD31, grey), neutrophils (Ly6G, purple) and CD45 (cyan). 8 minutes timelapse series were acquired, and neutrophils were tracked using Imaris software.

**Video 4: CD18 modulates lung neutrophil motility in tumour bearing mice.**

Lungs from 10-15 mm KP tumour bearing Catchup^IVM-red^ mice were harvested for live *ex vivo* multiplexed confocal microscopy of PCLS and stimulated *ex vivo* with either isotype control antibody or CD18 activating antibody (M18/2). Slices were stained for DNA (Hoechst, blue), vasculature (CD31, grey), neutrophils (Ly6G-td Tomato, purple), T-cells (CD8, green), alveolar macrophages (CD11c, orange) and leukocytes (CD45, cyan). 30 minutes timelapse series were acquired, and neutrophils were tracked using Imaris software.

**Video 5: G-CSF treatment modulates lung neutrophil motility in a CD18 dependent manner.**

Lungs from G-CSF treated mice were harvested for live *ex vivo* multiplexed confocal microscopy of PCLS and stimulated *ex vivo* with either isotype control antibody or CD18 activating antibody (M18/2). Slices were stained for DNA (Hoechst, blue), vasculature (CD31, grey) and neutrophils (Ly6G, purple). 15 minutes timelapse series were acquired, and neutrophils were tracked using Imaris software.

**Figure S1:**
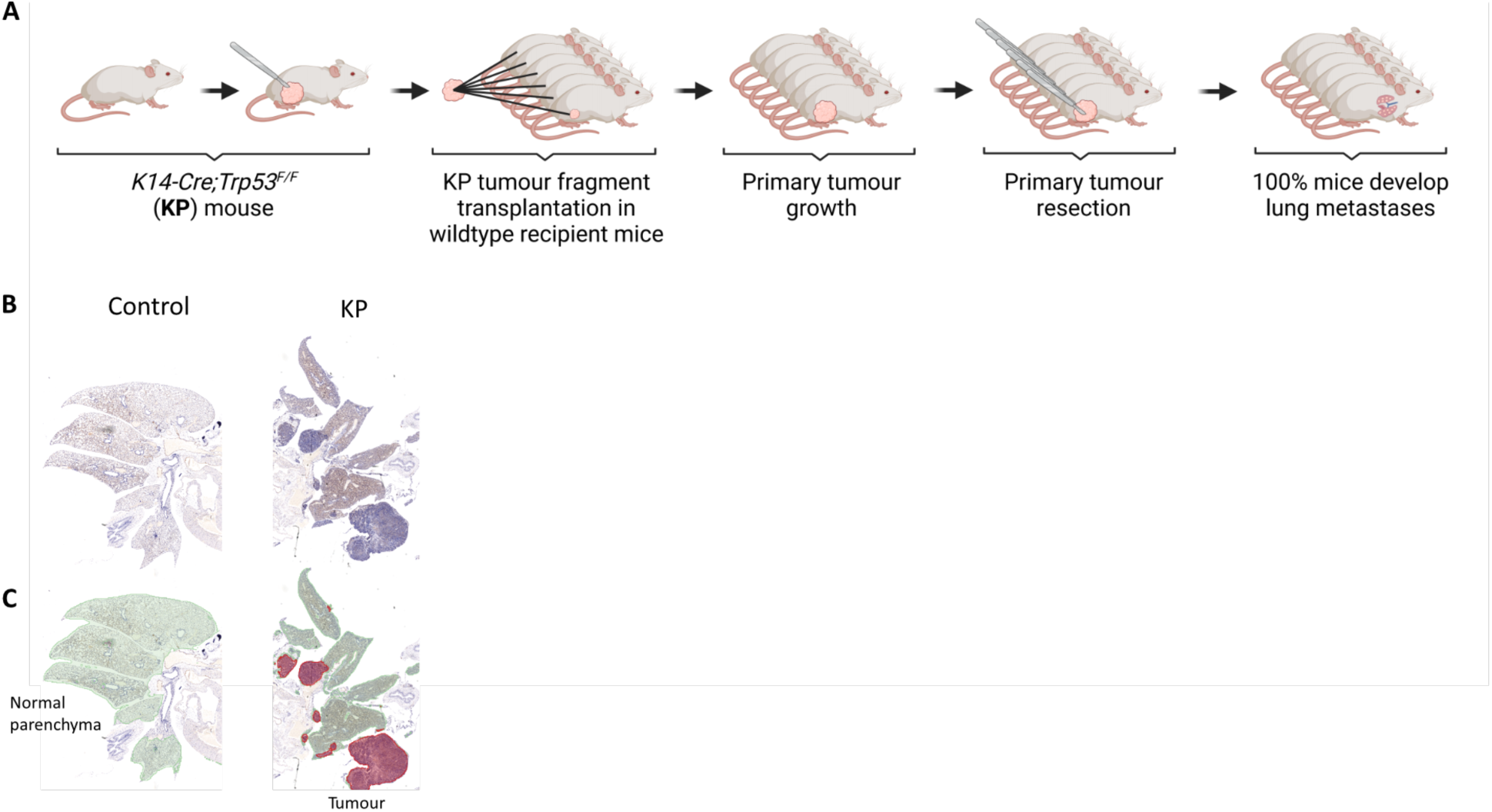
Spontaneous metastasis model. A) Mammary tumour fragments from KP mice were orthotopically transplanted into WT recipient mice, allowed to grow, then surgically resected when tumour reached 10 mm in any direction. Metastases develop in 100% of recipient mice. B) Representative hematoxylin & eosin staining from non-transplanted control and metastatic lungs. C) Annotation showing normal parenchyma (green) and tumour tissue (red) from a pixel classification in QuPath^45^.

**Figure S2:**
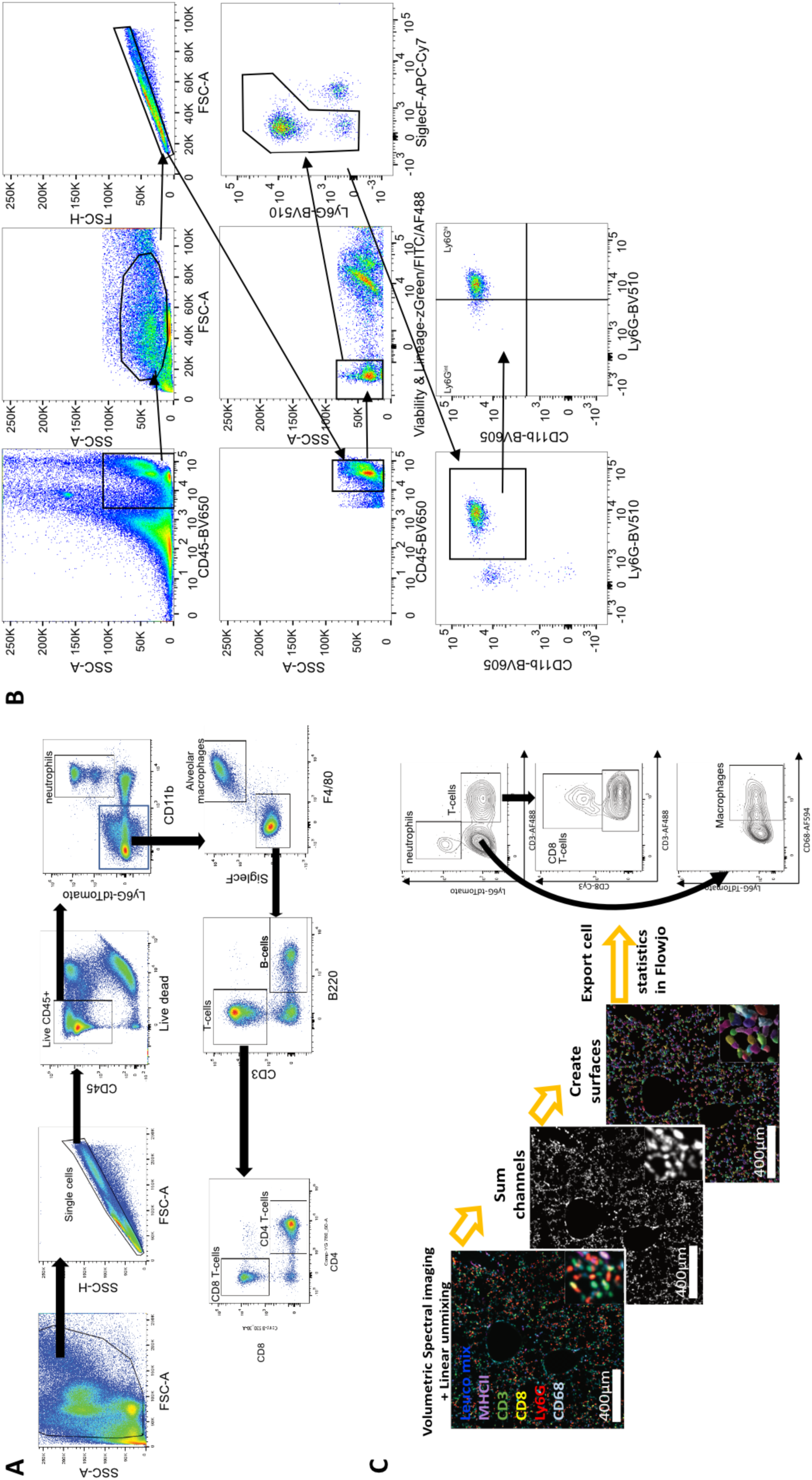
Gating strategies for flow cytometry and Histocytometry. A) Gating strategy for flow cytometry analysis of dissociated lung to phenotype CD45+ immune cell populations using lung immunophenotyping panel: Neutrophils (CD11b+ Ly6G+), Alveolar macrophages (CD11b-SiglecF+), B-cells (B220+), CD8 T-cells (CD3+ CD8+) and CD4 T-cells (CD3+ CD4+). Relevant to Figure 1. B) Gating strategy for flow cytometry analysis of Ly6G^int^ and Ly6G^hi^ neutrophil numbers, CD18, CD62L and CD101 neutrophil surface expression in Figures 1, 2 and 3 using neutrophil phenotyping panel. C) Histocytometry strategy for analysing lung tissue images in Figure 1E-G (Data shown here is the same image from Figure 1E for illustrative purposes). Fixed precision cut lung slices (PCLS) were stained with fluorescently labelled antibodies and imaged with a spectral confocal microscope. After linear unmixing of the different fluorophores, the channels corresponding to the different immune markers were summed to allow the segmentation of all immune cells in Imaris software. Cell statistics were then imported into FlowJo for analysis.

**Figure S3:**
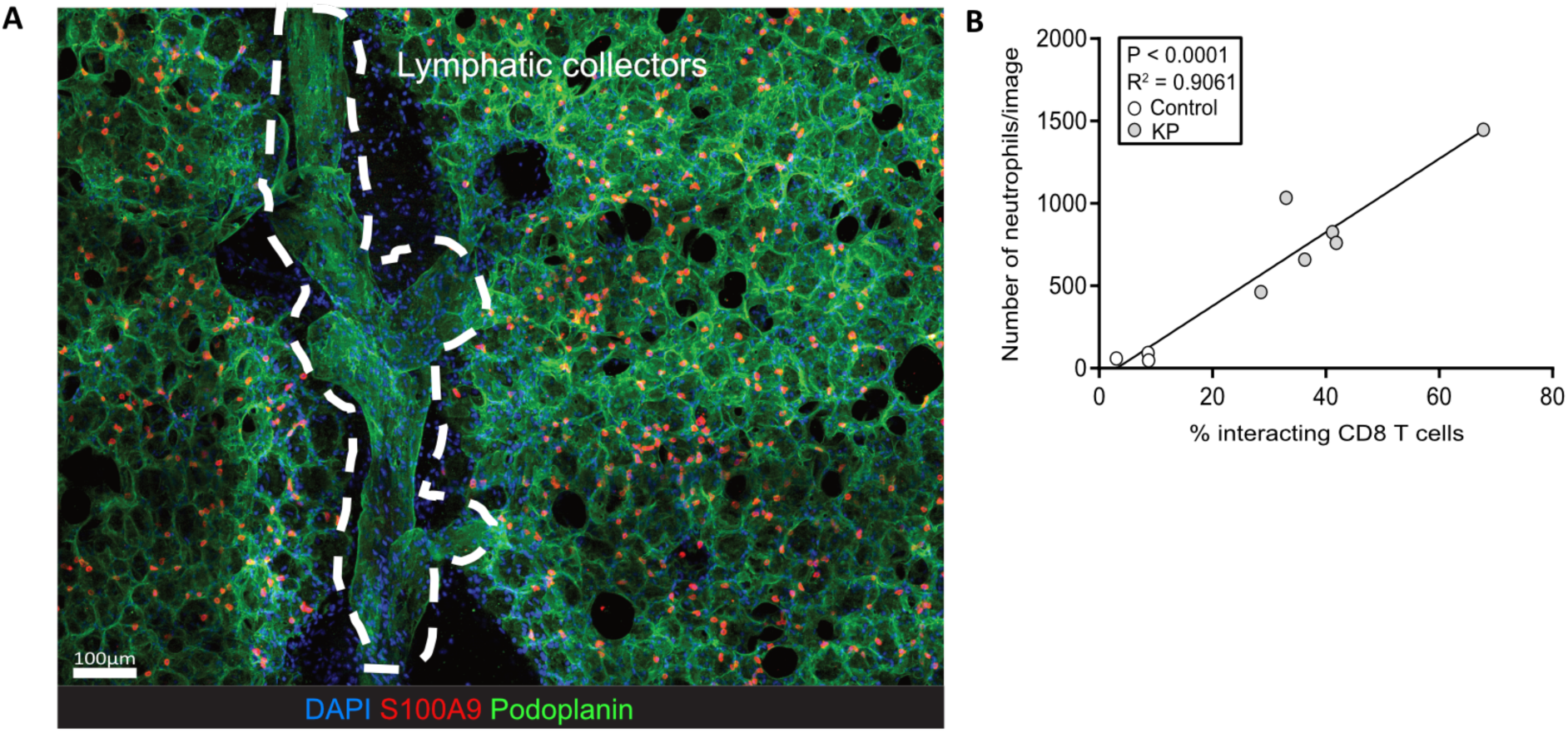
Localisation of lung neutrophils and CD8 T-cells interactions. (A) Podoplanin (green) staining for lymphatic collectors (it also stains alveolar cells in the lung) in a PCLS from a KP tumour bearing FVB/N mouse. S100A9 (red) was used to stain neutrophils and DAPI (blue) was used to stain DNA. (B) The correlation between CD8^+^ T cell:neutrophil interaction and neutrophil numbers was analysed using the Spearman’s rank correlation test (r = 0.8333 p=0.0083); then a linear regression was performed to describe relationship between variables, showing that 90% (R^2^= 0,9061, p<0.0001) of the interactions can be significantly explained by the number of neutrophils in the lung.

**Figure S4:**
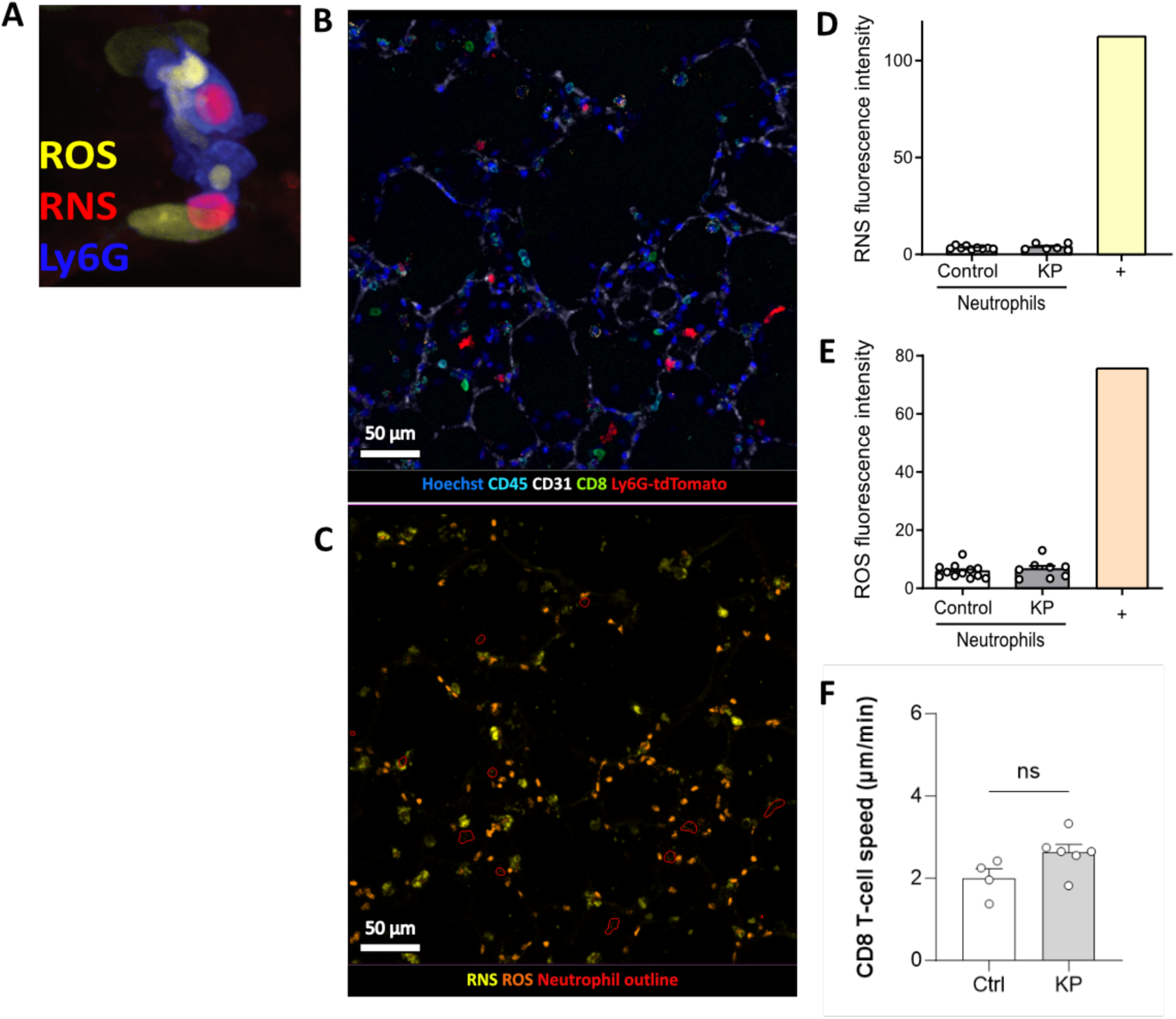
ROS/RNS production in live PCLS and CD8 T-cell speed. Lungs from control and KP tumour bearing mice were harvested for live *ex vivo* multiplexed confocal microscopy of PCLS to analyse their behaviour and free radical production (Reactive Oxygen Species – ROS and Reactive Nitrogen Species – RNS). (A) Positive control from a Phorbol 12-myristate 13-acetate (PMA) treated PCLS showing a neutrophil (Ly6G, blue) with cytoplasmic ROS (yellow) and nuclear RNS (red). (B-C) Slices were stained for DNA (Hoechst, blue), vasculature (CD31, grey), CD8 T-cells (CD8, green), leukocytes (CD45, cyan) and neutrophils (Ly6G, red) as well as probes for ROS (orange) and RNS (yellow). (D) Quantification of RNS mean fluorescence intensity in neutrophils. The + bar represents the mean intensity from a surrounding RNS positive cell. (E) Quantification of ROS mean fluorescence intensity in neutrophils. The + bar represents the mean intensity from a surrounding ROS positive cell. (D-E) Results are expressed as mean ± SEM of n = 8-12 mice per group. (F) CD8 T-cells mean speed in Control and KP orthotopic tumour-bearing mouse lung slices. Results are expressed as mean ± SEM of n = 4 control mice and 6 KP-bearing mice. A parametric T-test was performed, ns p>0.05.

**Figure S5:**
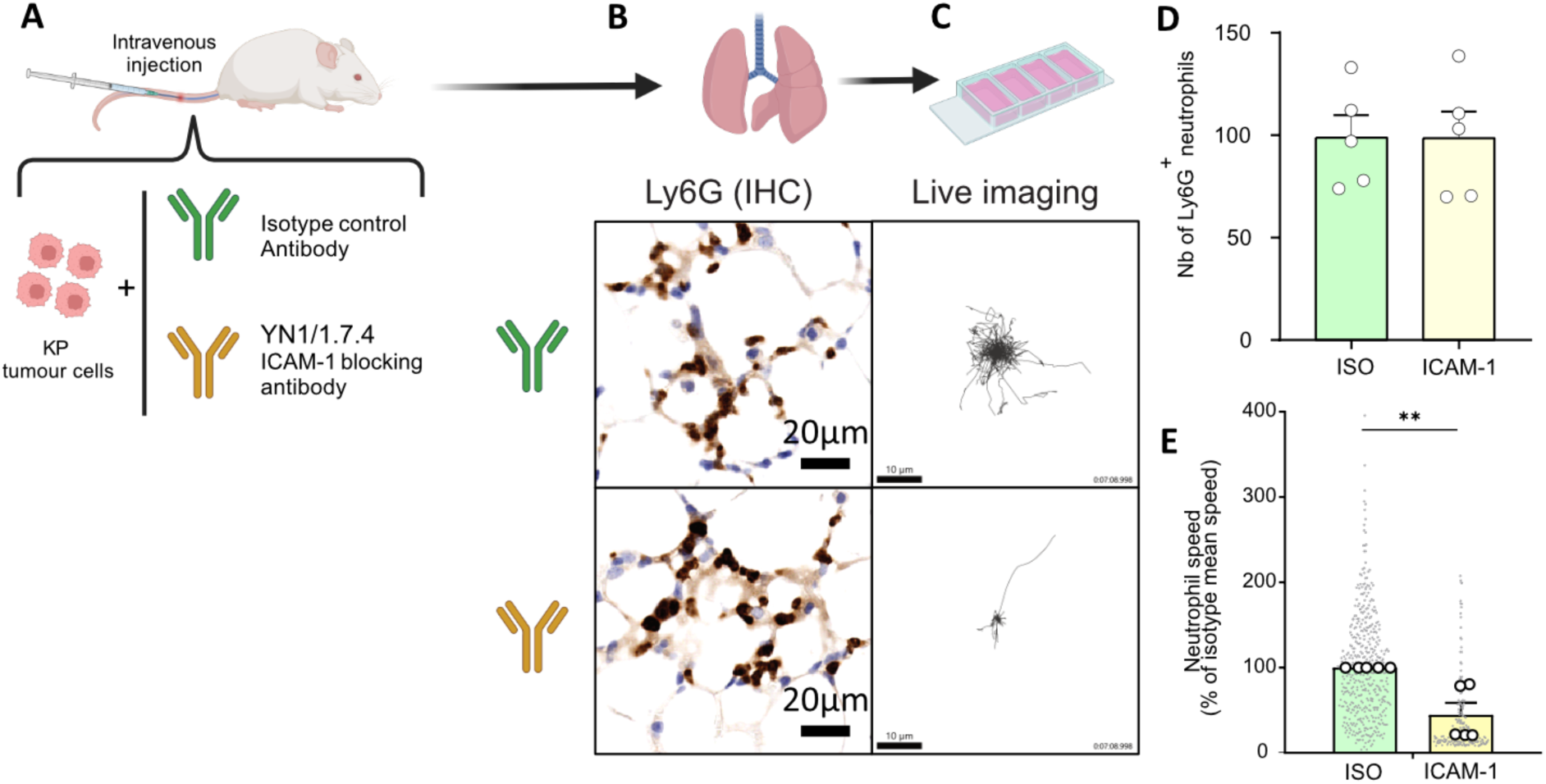
Modulation of neutrophil motility in an intravenous (IV) transplant model of KP cells, using ICAM-1 blocking antibodies. (A) Schematic of the experiment design. Mice were IV injected with KP cells and either isotype control antibody (green) or ICAM-1 blocking antibodies (yellow). 30 minutes later lungs were taken for live *ex vivo* confocal microscopy of PCLS (see video S3) and Immunohistochemistry. (B) Representative IHC images from the two treatment groups. (C) Representative neutrophil tracking plots from the two treatment groups. (D) Quantification of the number of neutrophils/mm^2^ in lung IHC images. Results are expressed as mean ± SEM of 5 mice per group. A T-test was performed. (E) Quantification of neutrophil speed as a % of the isotype control mean speed from live *ex vivo* microscopy. Grey dots represent the speed of each individual neutrophil (a total of 213-459 tracks per group were analysed) while white dots represent the mean neutrophil speed per mouse. Statistical analysis (nonparametric Mann-Whitney test) was performed on the mean value per animal. Results are expressed as mean ± SEM of n = 5 mice per group. **p≤0.01.

**Figure S6:**
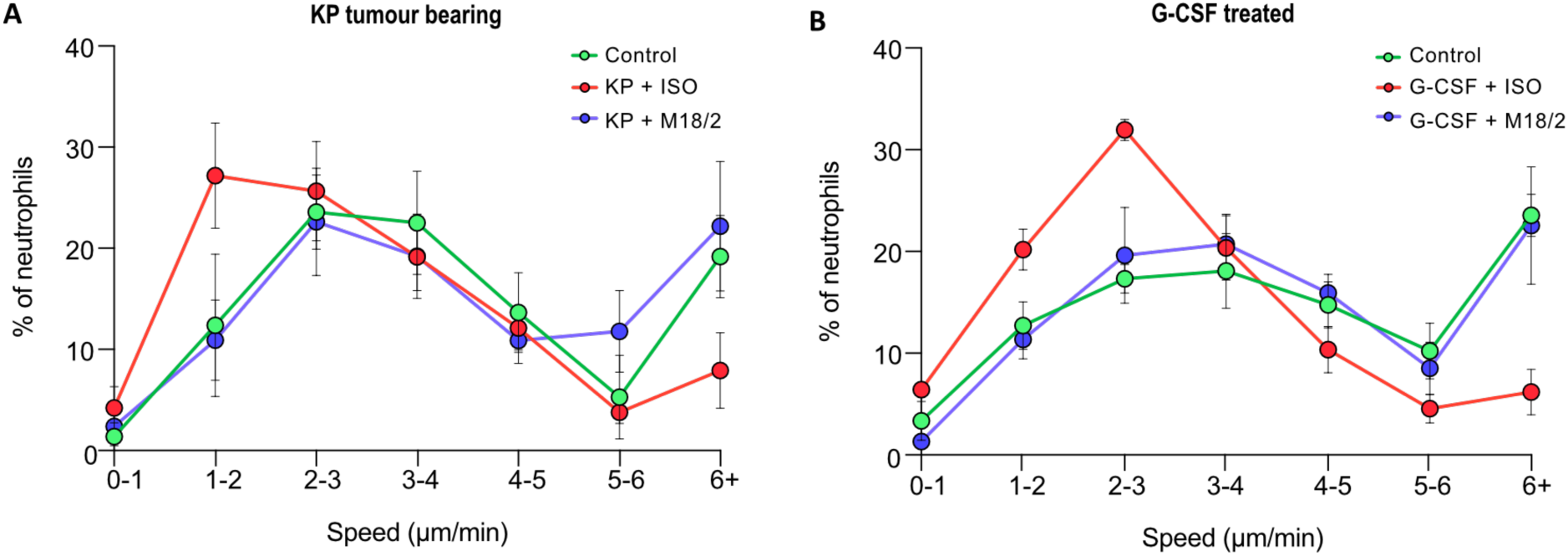
Neutrophil speed distribution in the lungs of mice with KP tumour and mice treated with G-CSF. Neutrophils were tracked in live multiplexed confocal microscopy timelapse images of *ex vivo* PCLS after *ex vivo* stimulation with either isotype control antibody or M18/2. (A) Speed distribution of neutrophils from control and KP tumour-bearing mice from Figure 2M. Results are expressed as mean +/-SEM of n = 4 control mice and 6 KP-tumour bearing mice (B) Speed distribution of neutrophils from PBS and G-CSF treated mice from Figure 3E. Results are expressed as mean +/-SEM of n = 4 PBS control and 5 G-CSF treated mice.

## Notes

### Summary of Updates

We have added additional data on CD62L expression (in Figure 3) and refined our description and interpretation in the text.

