## supplementary figures and legends for "Integrin conformation-dependent neutrophil slowing obstructs the capillaries of the pre-metastatic lung in a model of breast cancer"

**Video 5: G-CSF treatment modulates lung neutrophil motility in a CD18 dependent manner.**

Lungs from G-CSF treated mice were harvested for live *ex vivo* multiplexed confocal microscopy of PCLS and stimulated *ex vivo* with either isotype control antibody or CD18 activating antibody (M18/2). Slices were stained for DNA (Hoechst, blue), vasculature (CD31, grey) and neutrophils (Ly6G, purple). 15 minutes timelapse series were acquired, and neutrophils were tracked using Imaris software.

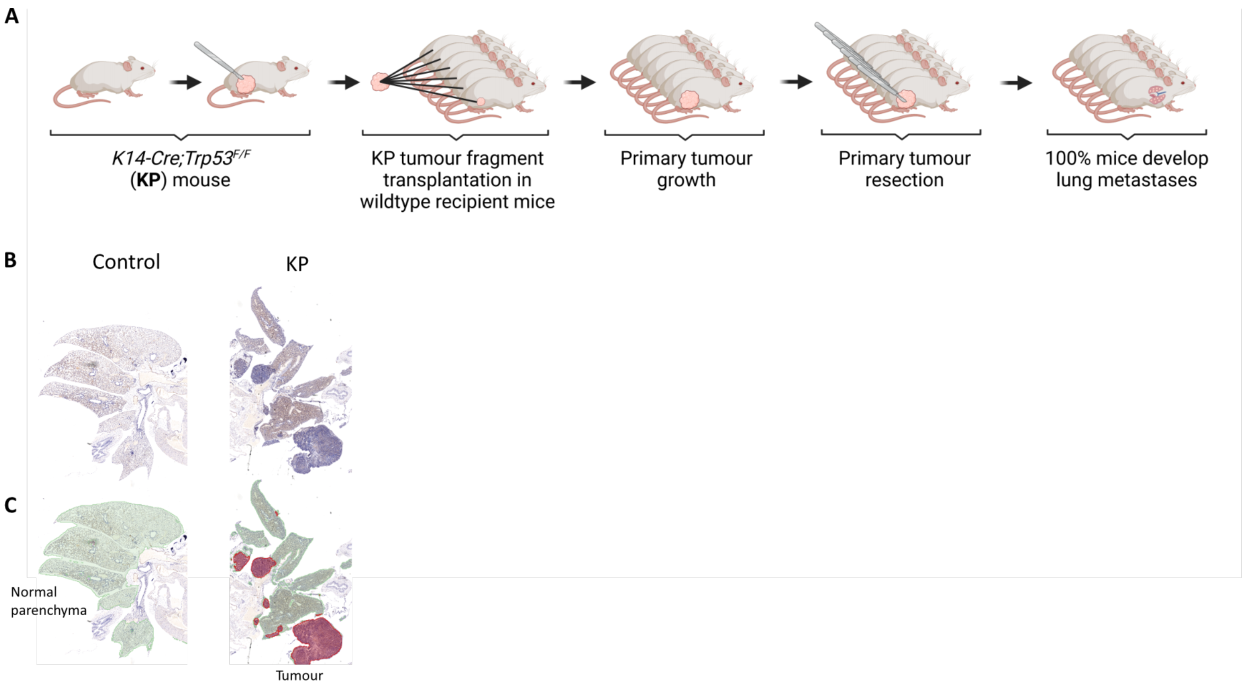


**Figure S1: Spontaneous metastasis model.**

A) Mammary tumour fragments from KP mice were orthotopically transplanted into WT recipient mice, allowed to grow, then surgically resected when tumour reached 10 mm in any direction. Metastases develop in 100% of recipient mice.

B) Representative hematoxylin & eosin staining from non-transplanted control and metastatic lungs.

C) Annotation showing normal parenchyma (green) and tumour tissue (red) from a pixel classification in QuPath[45].

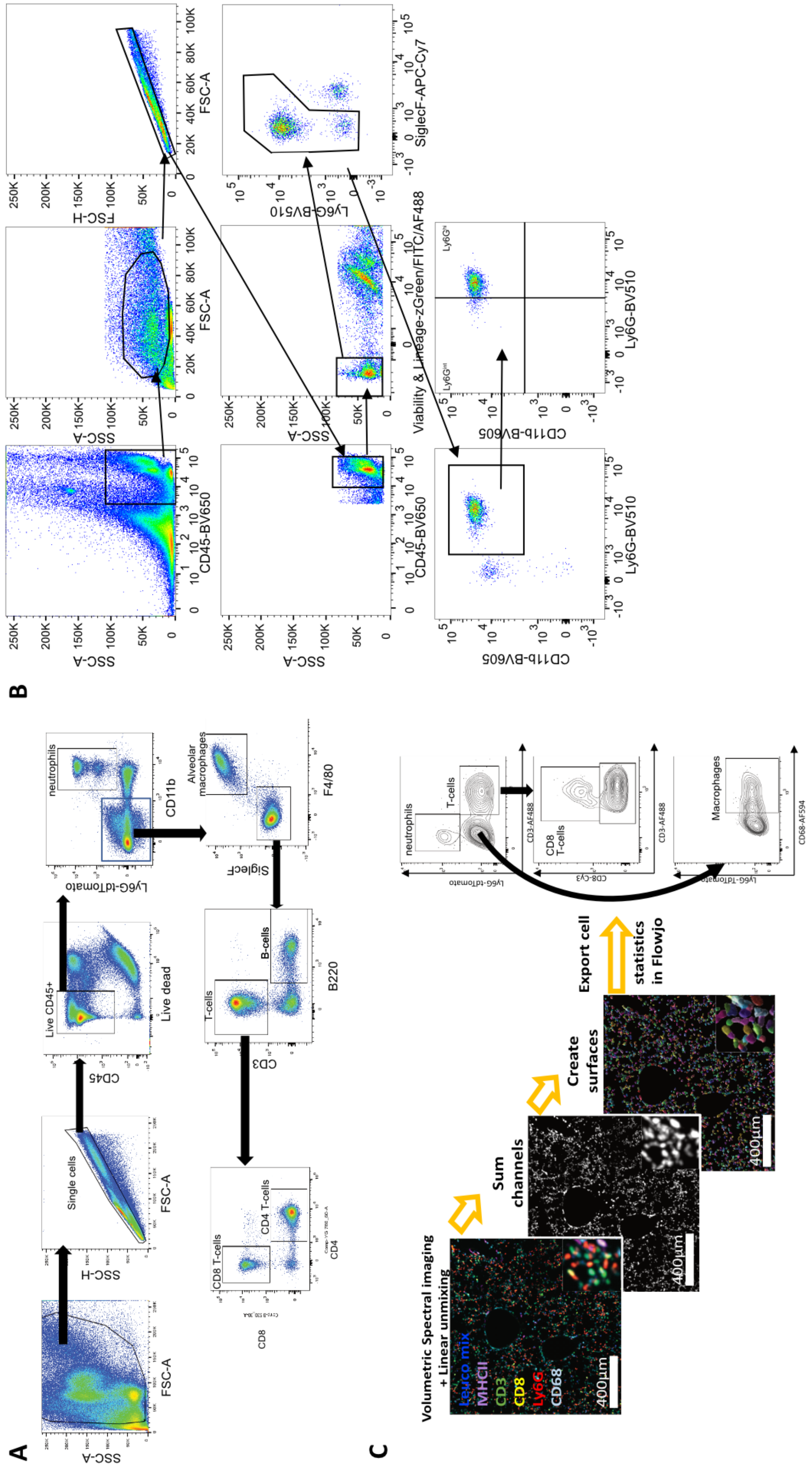


**Figure S2: Gating strategies for flow cytometry and Histocytometry.**

A) Gating strategy for flow cytometry analysis of dissociated lung to phenotype CD45+ immune cell populations using lung immunophenotyping panel: Neutrophils (CD11b+ Ly6G+), Alveolar macrophages (CD11b- SiglecF+), B- cells (B220+), CD8 T-cells (CD3+ CD8+) and CD4 T-cells (CD3+ CD4+). Relevant to Figure 1.

B) Gating strategy for flow cytometry analysis of $Ly6G^{int}$ and $Ly6G^{hi}$ neutrophil numbers, CD18, CD62L and CD101 neutrophil surface expression in Figures 1, 2 and 3 using neutrophil phenotyping panel.

C) Histocytometry strategy for analysing lung tissue images in Figure 1E-G (Data shown here is the same image from Figure 1E for illustrative purposes). Fixed precision cut lung slices (PCLS) were stained with fluorescently labelled antibodies and imaged with a spectral confocal microscope. After linear unmixing of the different fluorophores, the channels corresponding to the different immune markers were summed to allow the segmentation of all immune cells in Imaris software. Cell statistics were then imported into FlowJo for analysis.

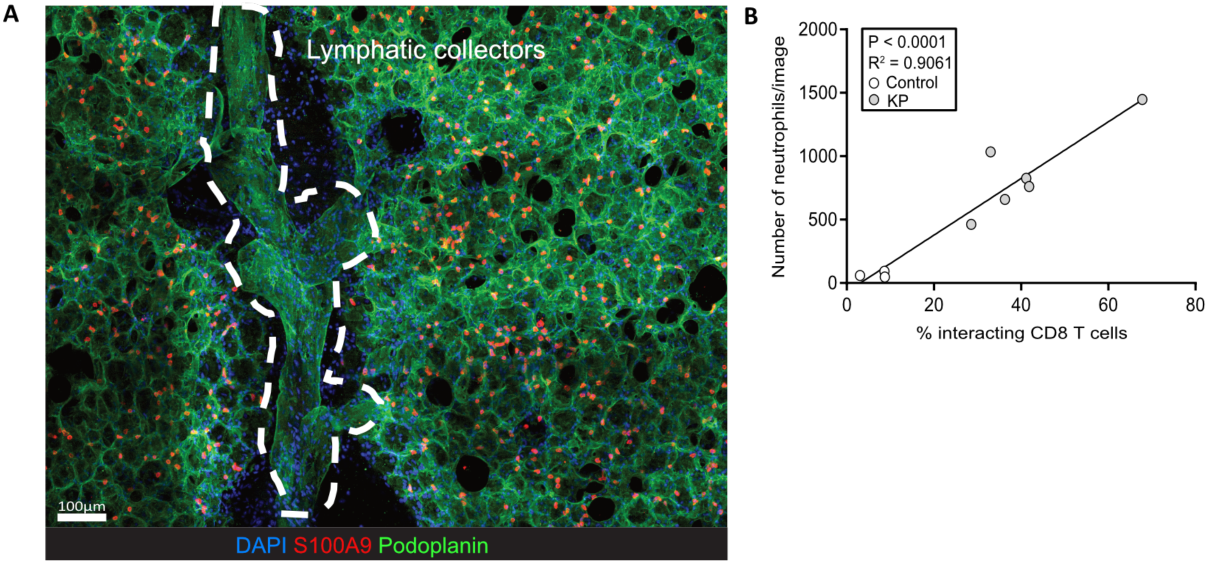


**Figure S3: Localisation of lung neutrophils and CD8 T-cells interactions.**

(A) Podoplanin (green) staining for lymphatic collectors (it also stains alveolar cells in the lung) in a PCLS from a KP tumour bearing FVB/N mouse. S100A9 (red) was used to stain neutrophils and DAPI (blue) was used to stain DNA.

(B) The correlation between $CD8^+$ T cell:neutrophil interaction and neutrophil numbers was analysed using the Spearman's rank correlation test ($r = 0.8333$ $p=0.0083$); then a linear regression was performed to describe relationship between variables, showing that 90% ($R^2= 0,9061$, $p<0.0001$) of the interactions can be significantly explained by the number of neutrophils in the lung.

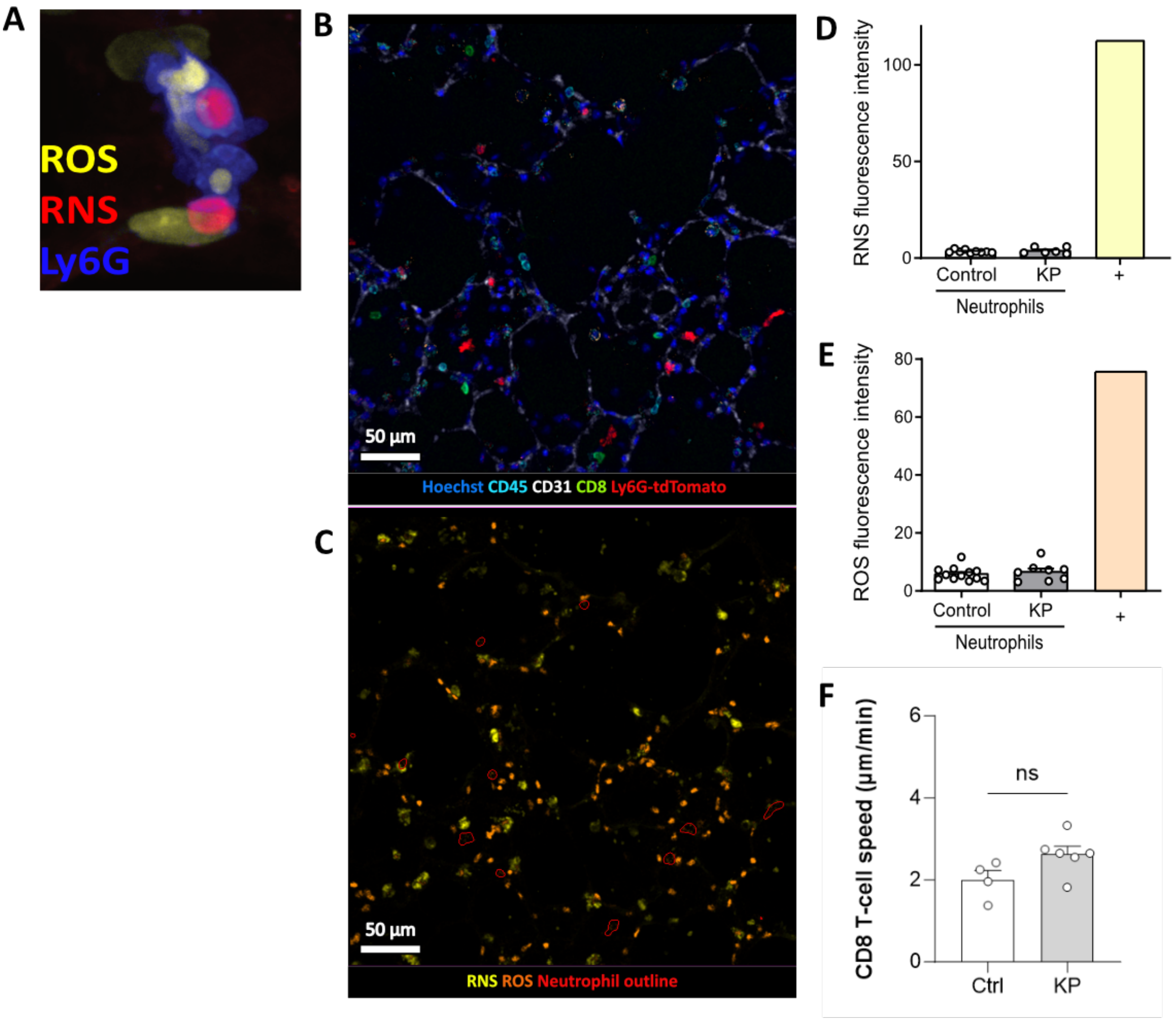


**Figure S4: ROS/RNS production in live PCLS and CD8 T-cell speed.**

Lungs from control and KP tumour bearing mice were harvested for live *ex vivo* multiplexed confocal microscopy of PCLS to analyse their behaviour and free radical production (Reactive Oxygen Species – ROS and Reactive Nitrogen Species – RNS).

(A) Positive control from a Phorbol 12-myristate 13- acetate (PMA) treated PCLS showing a neutrophil (Ly6G, blue) with cytoplasmic ROS (yellow) and nuclear RNS (red).

(B-C) Slices were stained for DNA (Hoechst, blue), vasculature (CD31, grey), CD8 T-cells (CD8, green), leukocytes (CD45, cyan) and neutrophils (Ly6G, red) as well as probes for ROS (orange) and RNS (yellow).

(D) Quantification of RNS mean fluorescence intensity in neutrophils. The + bar represents the mean intensity from a surrounding RNS positive cell.

(E) Quantification of ROS mean fluorescence intensity in neutrophils. The + bar represents the mean intensity from a surrounding ROS positive cell.

(D-E) Results are expressed as mean ± SEM of n = 8-12 mice per group.

(F) CD8 T-cells mean speed in Control and KP orthotopic tumour-bearing mouse lung slices. Results are expressed as mean ± SEM of n = 4 control mice and 6 KP-bearing mice. A parametric T-test was performed, ns $p>0.05$.

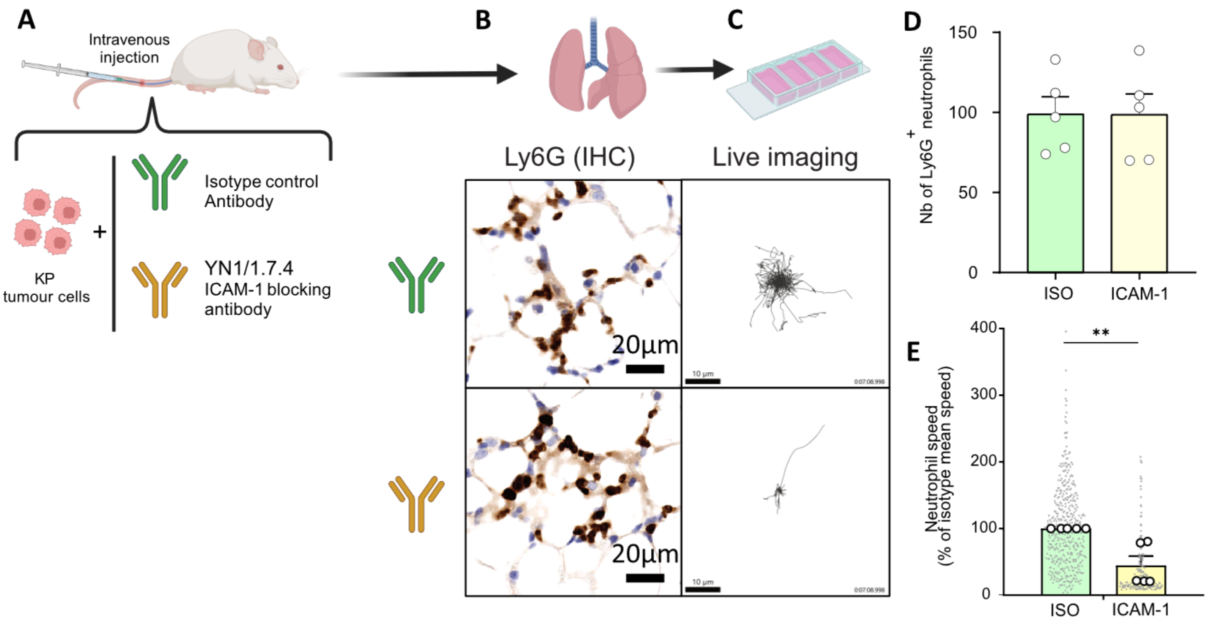


**Figure S5: Modulation of neutrophil motility in an intravenous (IV) transplant model of KP cells, using ICAM-1 blocking antibodies.**

(A) Schematic of the experiment design. Mice were IV injected with KP cells and either isotype control antibody (green) or ICAM-1 blocking antibodies (yellow). 30 minutes later lungs were taken for live *ex vivo* confocal microscopy of PCLS (see video S3) and Immunohistochemistry.

(B) Representative IHC images from the two treatment groups.

(C) Representative neutrophil tracking plots from the two treatment groups.

(D) Quantification of the number of neutrophils/$mm^2$ in lung IHC images. Results are expressed as mean ± SEM of 5 mice per group. A T-test was performed.

(E) Quantification of neutrophil speed as a % of the isotype control mean speed from live *ex vivo* microscopy. Grey dots represent the speed of each individual neutrophil (a total of 213-459 tracks per group were analysed) while white dots represent the mean neutrophil speed per mouse. Statistical analysis (nonparametric Mann-Whitney test) was performed on the mean value per animal. Results are expressed as mean ± SEM of n = 5 mice per group. **$p \leq 0.01$.

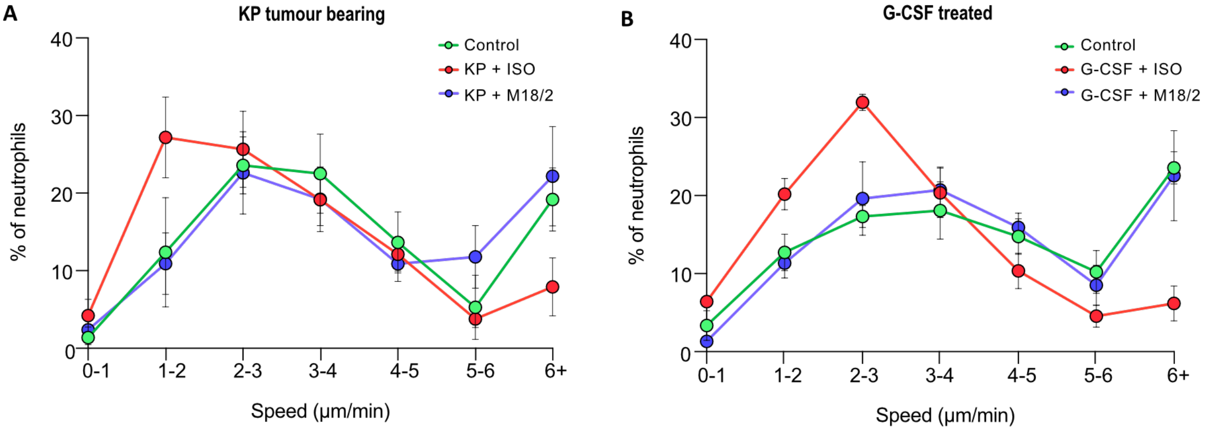


**Figure S6: Neutrophil speed distribution in the lungs of mice with KP tumour and mice treated with G-CSF.**

Neutrophils were tracked in live multiplexed confocal microscopy timelapse images of *ex vivo* PCLS after *ex vivo* stimulation with either isotype control antibody or M18/2.

(A) Speed distribution of neutrophils from control and KP tumour-bearing mice from Figure 2M. Results are expressed as mean +/- SEM of n = 4 control mice and 6 KP-tumour bearing mice

(B) Speed distribution of neutrophils from PBS and G-CSF treated mice from Figure 3E. Results are expressed as mean +/- SEM of n = 4 PBS control and 5 G-CSF treated mice.
